# How well does electrical source imaging reconstruct neural time series? Evidence from simultaneous scalp and intracranial EEG recordings

**DOI:** 10.64898/2026.10.26.761837

**Authors:** Maria Dimou, Steven Beumer, Andrea Pigorini, Ezequiel Mikulan, Alessandra Calcagno, Laurent Spinelli, Shahan Momjian, Margitta Seeck, Serge Vulliemoz, Nicolas Roehri

## Abstract

Electrical source imaging (ESI) estimates neural activity from scalp EEG, but its ability to reconstruct the temporal dynamics of spontaneous brain activity beyond source localization remains unclear. This is assessed using simultaneous high-density EEG and intracranial EEG (iEEG) recordings during spontaneous activity in 34 patients with drug-resistant focal epilepsy. Four ESI approaches (three distributed source models and one beamformer) are applied to estimate neural activity, which is then projected onto iEEG electrode locations to generate virtual iEEG signals for comparison with measured recordings. Reconstruction fidelity is quantified using Pearson correlations between actual and virtual iEEG signals, while spatial accuracy is assessed through peak localization error and spatial deviation derived from point-spread and cross-talk functions. Analyses are performed in broadband and delta, theta, alpha, and beta bands. Temporal reconstruction remains limited, with the highest estimated correlation reaching 0.126, localization errors of approximately 3 cm, and substantial spatial leakage. Reconstruction quality decreases with electrode depth and increasing frequency, while improving with higher epileptic spike rates. No single method is optimal across all metrics, although beamforming shows the weakest temporal reconstruction. These findings indicate that ESI only partially captures spontaneous intracranial dynamics and define spatial and temporal constraints for interpreting EEG- based source reconstructions.

## 1. Introduction

Electroencephalography is a non-invasive tool with millisecond temporal resolution that is particularly valuable for studying rapidly evolving neural time series. This is especially relevant in disorders such as epilepsy, where pathological activity evolves dynamically over time (Lisgaras et al. 2025). In studies of both healthy and pathological brain function, characterizing the temporal dynamics and interactions between brain regions is central to many neuroscientific investigations. Interpreting scalp EEG in terms of these dynamics requires source reconstruction to accurately recover the temporal evolution of neural activity, an assumption that remains insufficiently validated.

EEG signals recorded at the scalp reflect a mixture of electrical activity generated by multiple neural sources within the brain. As different configurations of neural sources can produce identical scalp signals, estimating the location and activity of the underlying brain sources–known as the EEG inverse problem–is inherently ill-posed (Phillips et al. 2026). To address this issue, electrical source imaging (ESI) approaches attempt to estimate the underlying neural activity by introducing biophysical and mathematical constraints that restrict the set of possible solutions (Michel and Brunet 2019). Different choices of these constraints have led to the development of a variety of ESI approaches (Grech et al. 2008).

Among the most widely used ESI approaches are distributed source models, particularly minimum- norm methods. These compute neural activity by imposing L2-norm regularization, favoring solutions with minimal overall source power, and include weighted minimum-norm estimates (wMNE), standardized low-resolution brain electromagnetic tomography (sLORETA), and exact low-resolution brain electromagnetic tomography (eLORETA) (Lin et al. 2006; Pascual-Marqui 2002, 2007). Each incorporates weighting or normalization to reduce the inherent bias toward superficial sources. Alternative approaches based on L1-norm (Uutela et al. 1999; Ding and He 2008) or mixed norm regularization (Gramfort et al. 2012) promote sparse source configurations, encouraging solutions with a limited number of sources and thus providing a compromise between focal dipole models and fully distributed solutions. However, these methods are computationally demanding and may produce temporally discontinuous source estimates, limiting their suitability for the analysis of continuous time series. A separate class of methods uses spatial filtering. For example, linearly constrained minimum variance (LCMV) beamformers (Van Veen et al. 1997) estimate activity at a given location by constructing adaptive spatial filters that suppress signals from other locations, although their performance may be affected by temporal correlation between sources. These approaches differ in how they constrain the solution and have been compared and characterized in several studies. Comparisons have focused primarily on their localization properties (Hauk et al. 2022; Pascarella et al. 2023), that is, how accurately and focally they estimate the spatial origin of neural activity. However, validation of ESI is challenging because the true sources of neural activity are generally unknown.

Existing studies have addressed this challenge using different validation strategies. One approach relies on simulated source data, for which the ground truth is known (Samuelsson et al. 2021), although the generalizability of these results to real-world recordings may be limited. Another approach uses an indirect clinical reference by comparing the location of EEG source-localized epileptic spikes to the resection area in patients with epilepsy who achieved favorable surgical outcomes (van Mierlo et al. 2017; Spinelli et al. 2023). Intracranial EEG (iEEG) recordings from depth electrodes implanted during presurgical evaluation of patients with drug-resistant epilepsy provide direct measurements of neural activity at specific brain locations. Although their spatial coverage is limited by clinical considerations, iEEG recorded simultaneously with scalp EEG offers a local electrophysiological reference for evaluating EEG source estimates. These recordings have recently been used for a more direct form of validation during electrical stimulation (Mikulan et al. 2020; Pascarella et al. 2023).

Nevertheless, previous validation studies have focused primarily on source localization accuracy, evaluating whether reconstructed sources correctly identify the spatial origin of neural activity.

Although these studies provide important insight into the spatial localization properties of ESI methods, to our knowledge no previous study has investigated whether ESI can accurately reconstruct the temporal evolution of spontaneous neural activity. Such reconstruction presents an additional challenge because spontaneous neural activity may have a lower signal-to-noise ratio (SNR) than evoked activity (e.g., event-related potentials and electrically evoked responses), and its temporal fidelity can only be evaluated using simultaneous iEEG and scalp EEG recordings.

In this study, we investigated how accurately different ESI methods capture the temporal dynamics of neural activity using a rare dataset of simultaneous scalp hdEEG and intracranial EEG acquired during spontaneous activity in quiet wakefulness. This dataset provides a unique opportunity to validate EEG source estimates directly against intracranial recordings at the implanted electrode locations. We evaluated the performance of wMNE, sLORETA, eLORETA, and the LCMV beamformer using measures of temporal fidelity and spatial accuracy, including localization error and spatial spread. We then compared the methods based on these metrics and investigated how their performance varied across frequency bands and with electrode depth and spike rate.

## 2. Materials and methods

### 2.1 Cohort

We retrospectively analyzed data from patients with drug-resistant focal epilepsy that were evaluated for epilepsy surgery at two different centers: the Epilepsy Unit of University Hospitals of Geneva, Switzerland, and the Niguarda Hospital of Milan, Italy. We included patients with (i) focal epilepsy, (ii) simultaneous hdEEG (256 electrodes) and iEEG data, (iii) magnetic resonance imaging (MRI) scans acquired before the iEEG implantation, and (iv) computed tomography (CT) scans acquired after the implantation. Data from patients with poor EEG recording quality were excluded. Patient demographic and clinical characteristics are summarized in Table S1. Detailed descriptions of the data acquisitions are provided in the corresponding papers (De Stefano et al. 2021; Roehri et al. 2026; Mikulan et al. 2020).

### 2.2 Preprocessing

#### 2.2.1 Segmentation and parcellation

T1w MRIs were used for brain segmentation and parcellation into regions of interest (ROIs) using Freesurfer (Fischl 2012), based on the Desikan-Killiany cortical atlas. We then constructed a gray matter mask atlas to define the source space. Among the gray matter regions, we included the hippocampus and amygdala, as commonly done in epilepsy studies, but excluded the thalamus, caudate, putamen, pallidum, and accumbens area.

#### 2.2.2 iEEG localization

MRIs and CTs were linearly co-registered and used for the localization of the iEEG electrodes using Cartool software (Brunet et al. 2011) for the data of University Hospitals of Geneva, while the procedure for the Niguarda Hospital (Milan) dataset has been described previously (Mikulan et al. 2020). Each iEEG electrode was assigned to gray or white matter by overlaying its coordinates onto the gray matter mask atlas.

#### 2.2.3 hdEEG and iEEG

First, we applied a zero-phase 4th-order Butterworth bandpass filter between 1 and 45 Hz and a bandstop filter between 48 and 52 Hz to attenuate the power line noise, and then downsampled the hdEEG and iEEG signals to 200 Hz. For hdEEG, we performed independent component analysis to identify and reject components reflecting cardiac, oculomotor, and muscle activity (Jung et al. 2000). For iEEG, only electrodes within the gray matter were retained for the analysis, and we applied common average re-referencing computed using only gray-matter electrodes, thereby attenuating components shared across channels, including the contribution of the original recording reference. Preprocessing was performed using custom MATLAB scripts based on the FieldTrip toolbox (Oostenveld et al. 2011), with default parameters unless otherwise stated. We analyzed broadband signals as well as narrowband signals corresponding to the canonical frequency bands, ranging from delta to beta.

### 2.3 Electrical source imaging

#### 2.3.1 ESI methods

For the forward problem, we computed a three-layer head model (scalp, skull, and brain), using the Boundary Element Method implemented in OpenMEEG (Gramfort et al. 2010).

Dipoles with unconstrained orientations were uniformly distributed on a 6-mm grid throughout the gray matter across the entire brain, including the hippocampus and amygdala, while ensuring they were not placed within the thalamus and basal ganglia. Using the headmodel and the source model, we computed the leadfield matrix.

We then employed different ESI methods, i.e., eLORETA, sLORETA, wMNE and the LCMV beamformer to reconstruct the brain activity at the source level. Source reconstruction was performed using custom MATLAB scripts utilizing the FieldTrip toolbox. To ensure a fair comparison across methods, λ was set to 5%. Details of the ESI methods are provided in the Supporting Information.

### 2.4 Virtual iEEG electrodes and performance assessment

Comparing the source-reconstructed EEG signals directly with the iEEG signals poses a challenge: the reconstructed sources are distributed throughout the brain and do not necessarily coincide with the iEEG electrode positions.

#### 2.4.1 Virtual iEEG electrodes

To enable a direct comparison, we employed an infinite volume conductor forward model that calculates, at each iEEG electrode location, the electric potential created by a current dipole (Grova et al. 2016). Specifically, assuming an electrical conductivity *σ*, the potential at iEEG electrode j, located at position *E*_j_, generated by a unit current dipole at source location *S_i_*, with orientation *n→_i_* is given by:

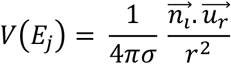

where *u_r_* is the unit vector pointing from *S_i_* to *E*_j_, and *r* is the Euclidean distance between *S_i_* and *E*_j_. We used the same parameter values as in previous work (Grova et al. 2016; Afnan et al. 2025).

Using this model, we computed the electric potentials that the activity of the current dipoles would produce at the positions of the implanted iEEG electrodes. This projection enables a one-to-one comparison with the iEEG channels, and we refer to the projected signals as virtual iEEG. The virtual iEEG signals were not re-referenced, since they are biased estimates, affected by spatial leakage and therefore highly correlated with each other. Applying the same common average re-referencing as for the iEEG signals would mix them further.

#### 2.4.2 iEEG-virtual iEEG correlation

To quantify how well the time series of the virtual iEEG were reconstructed by each ESI method, we computed the correlation matrix between iEEG and virtual iEEG using Pearson correlation. We chose the correlation matrix over the covariance matrix to focus on temporal fidelity, independently of relative amplitude differences, as in functional connectivity measures. Since the polarity of iEEG signals is ambiguous, we used the absolute values of the correlation. First, we examined the diagonal of the matrix, which contained correlations for pairs of corresponding iEEG electrodes and virtual iEEG electrodes. Notably, this metric, which we refer to from now on as the iEEG-virtual iEEG correlation, or simply correlation, primarily assesses time-series reconstruction performance. In other words, it measures how well the signal recorded by each iEEG electrode is reconstructed at its own location. Subsequently, we used the entire correlation matrix to assess the spatial resolution of the different inverse solutions, as detailed in the following section. In addition, we assessed whether time-series reconstruction performance varied across frequency bands (broadband, delta, theta, alpha, and beta) by comparing the iEEG-virtual iEEG correlation for each ESI method. The comparison framework is illustrated in Figure 1.

**Figure 1.**
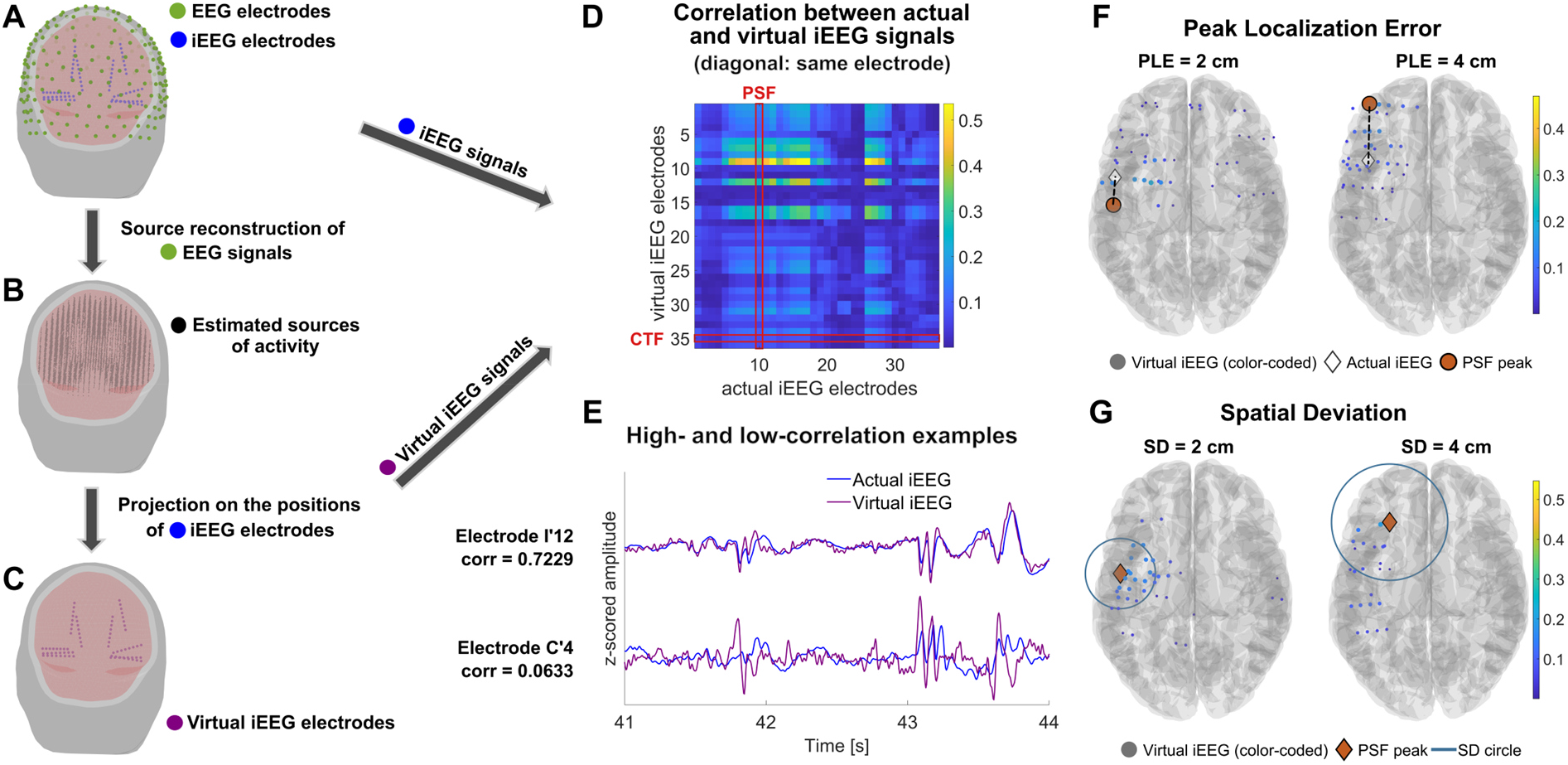
Framework for comparing virtual and actual iEEG signals at corresponding electrode locations, and definition of the reconstruction metrics. (A) Head model showing the scalp EEG electrodes (green dots) and the implanted iEEG electrodes (blue dots). The two modalities were recorded simultaneously. (B) Sources of brain activity (black dots) estimated through electrical source imaging (ESI). (C) Virtual iEEG electrodes (purple dots), obtained by projecting the source signals onto the iEEG electrode locations. (D) Absolute Pearson correlations between actual and virtual iEEG signals of a representative patient; each column is a point-spread function (PSF) and each row a cross-talk function (CTF). One of each is outlined in red, but all were used in the analysis. (E) Actual (blue) and virtual (purple) iEEG time series for two electrodes, with a high (top) and a low (bottom) diagonal correlation—i.e., between an actual electrode and its own virtual counterpart—reflecting good and poor reconstruction, respectively. (F) Examples of varying peak localization error (PLE): the distance between the PSF/CTF peak and the corresponding iEEG electrode location. (G) Examples of varying spatial deviation (SD): the spatial spread of the reconstruction around the PSF/CTF peak. In (F) and (G), each example shows a single PSF. Panels (D)–(G) show examples from different patients. In the following figures, PSF-PLE denotes the PLE of the PSF (similarly for PSF-SD, CTF-PLE and CTF-SD).

#### 2.4.3 Reconstruction characterization metrics

In our framework, we considered the signals recorded from the implanted iEEG electrodes to be the ground truth, and here we aimed to examine the localization accuracy and spatial spread of the estimated virtual iEEG signals. We interpreted the correlation matrix between iEEG and virtual iEEG signals as an analogue of the resolution matrix (Hauk et al. 2022), which maps the relationship between the sources and their estimates in classical ESI. This analogy allowed us to apply two well-established concepts of the resolution matrix—namely the point-spread function (PSF) and cross-talk function (CTF)—to our context. In the correlation matrix, each column can be interpreted as a PSF, illustrating the spread of activity from a single iEEG electrode across the virtual iEEG electrodes. Similarly, each row can be interpreted as a CTF, reflecting how activity from multiple iEEG electrodes contributes to a single virtual iEEG electrode.

To characterize our reconstruction, we employed two metrics based on either the PSF or CTF: peak localization error (PLE) and spatial deviation (SD). PLE quantifies the localization accuracy as the Euclidean distance between the actual location of the *i*-th iEEG electrode (*r^iEEG^*) and the location of the maximum value of its corresponding PSF/CTF (*r^p^*):

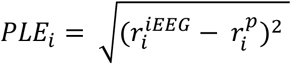

Ideally, the maximum PSF/CTF value of electrode *i* should coincide with the true location of electrode *i*, resulting in a PLE of 0 cm. In the correlation matrix, this would correspond to the maximum value occurring on the diagonal (Figure 1D). Specifically, PSF-PLE quantifies where a given iEEG electrode’s activity appears in the virtual reconstruction, while CTF-PLE shows which iEEG electrode is the dominant driver of a given virtual electrode’s reconstructed signal. It is worth noting that these two quantities are not reciprocal: the *i*-th iEEG electrode may be best reconstructed by a virtual electrode *j*, while the virtual electrode *i* may be driven most strongly by a different iEEG electrode *k*.

For each iEEG electrode *i*, SD measures the correlation-weighted spread of the virtual iEEG electrode locations around its PSF/CTF peak (Figure 1G) and is defined as:

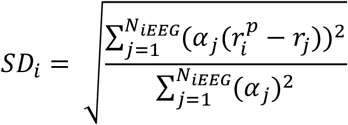

where *N_iEEG_* is the number of iEEG electrodes, *α*_j_ is the absolute correlation value of virtual electrode *j* with the *i*-th iEEG electrode, *r*_j_ is the location of electrode *j*, and *r^p^* is the location of the PSF/CTF peak for electrode *i*. Again, PSF-SD and CTF-SD address complementary questions: PSF-SD quantifies how much the reconstruction of a given iEEG electrode’s activity leaks around its PSF peak (i.e., the virtual iEEG electrode with the highest correlation), while CTF-SD quantifies how broadly a given virtual iEEG electrode is influenced by surrounding actual iEEG electrodes. All subsequent analyses described below were performed separately for the PSF- and CTF-based metrics. To verify that the reconstruction metrics behave as expected, we tested the hypothesis that lower PLE and SD values are associated with higher iEEG-virtual iEEG correlation.

#### 2.4.4 Impact of electrode depth and spike rate

We also examined the effect of iEEG electrode depth and iEEG spike rate on the reconstruction quality as measured by the iEEG-virtual iEEG correlation. We used Delphos software (Roehri et al. 2016, 2017) to automatically detect spikes in the iEEG electrode time series. The spike rate was computed as the number of detected spikes per minute for each iEEG channel. The electrode depth was computed as the Euclidean distance between the iEEG electrode’s position and the position of the nearest scalp electrode. Electrode depth was measured in cm, and spike rate was log-transformed as 10log4o(spikes/min). We repeated the analysis for all four ESI methods and all five frequency bands.

#### 2.4.5 Regions-of-interest analysis

To assess whether reconstruction accuracy varied across brain regions, we mapped each iEEG electrode to its corresponding brain area using the Lausanne 2018 parcellation atlas at scale 3 (231 regions) (Tourbier et al. 2022a, 2022b). We excluded bilateral basal ganglia structures (caudate, putamen, pallidum, nucleus accumbens), thalamus, and brainstem from the analysis, consistent with the source model used for EEG source reconstruction. This resulted in 220 regions included in the analysis (216 cortical parcels plus bilateral hippocampus and amygdala). Additionally, we mapped the left- hemisphere ROI labels onto their corresponding right-hemisphere ROI labels, pooling both hemispheres to increase statistical power. For each ROI, we estimated the iEEG-virtual iEEG correlation and reconstruction metrics (PLE and SD, for both PSFs and CTFs). We included only ROIs sampled by at least 8 patients to obtain more statistically robust results, resulting in 37 ROIs.

### 2.5 Statistical analysis

#### 2.5.1 Effects of frequency band and ESI method on source reconstruction accuracy

We fitted separate generalized linear mixed-effects (GLME) models to test the effects of frequency band and ESI method on each aspect of source reconstruction accuracy, namely the iEEG-virtual iEEG correlation and the PLE and SD from both PSFs and CTFs. Each model included main effects of frequency band and ESI method, as well as their interaction term. Random intercepts were included for patient and electrode nested within patient. We used a gamma distribution, which is suitable for positive and skewed data. For the absolute iEEG-virtual iEEG correlation, we used a logit link function to constrain the fitted values between 0 and 1, and for PLE and SD a log link function. The general model formula was:

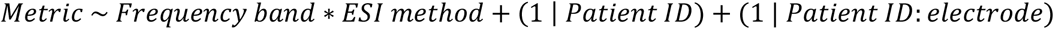

Model fit is reported as the adjusted R² throughout. We used ANOVA to assess the main effects and their interaction, and we conducted post hoc analysis to examine: (1) pairwise frequency band differences within each ESI method, and (2) pairwise ESI method differences within each frequency band. All post hoc pairwise comparisons were Bonferroni-corrected for multiple comparisons (10 comparisons for frequency bands within each method; 6 comparisons for methods within each frequency band). We reported the estimated marginal means back-transformed to the response scale, with 95% confidence intervals and corrected p-values. These are referred to as estimated values throughout the manuscript.

To aggregate performance, we ranked the four ESI methods for each frequency band using a Copeland- style scoring system (3 points when a method was significantly better in a pairwise comparison, 1 point when there was no significant difference, and 0 points when a method was significantly worse). We performed separate rankings for PSF-based metrics (combining iEEG-virtual iEEG correlation, PSF-PLE, and PSF-SD) and CTF-based metrics (combining iEEG-virtual iEEG correlation, CTF-PLE, and CTF-SD).

#### 2.5.2 Characterization of the reconstruction metrics

We fitted separate GLME models for each ESI method and frequency band to examine the relationship between the iEEG-virtual iEEG correlation and the reconstruction metrics. The model formulas were:

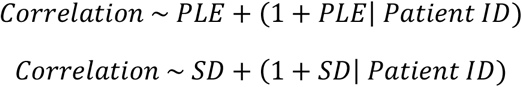

One model was fitted per combination of ESI method, frequency band, and reconstruction metric (i.e., PLE and SD for PSF and CTF).

We also examined the effect of electrode depth and spike rate on the iEEG-virtual iEEG correlation, using separate GLME models for each ESI method and frequency band. The model formulas were:

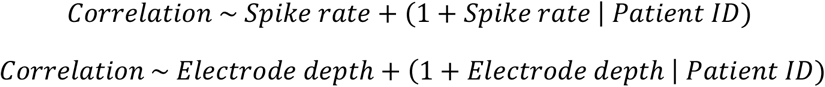

All models used a gamma distribution with a logit link function. The random effects accounted for patient-specific baseline correlation levels (random intercepts) and patient-specific relationships between each predictor and the correlation (random slopes). We computed 95% confidence intervals for fixed effects. Bonferroni correction was applied separately for each ESI method and each set of predictors (PSF-based PLE and SD; CTF-based PLE and SD; spike rate and electrode depth), across the five frequency bands and the two predictors of each set (5 × 2 = 10 comparisons per ESI method and predictor set).

#### 2.5.3 Effects of region of interest

To assess whether ROI influences the correlation or reconstruction metrics, we fitted separate GLME models for each frequency band and metric, with the general formula:

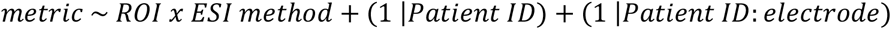

All models used a normal distribution with an identity link function. We included an ROI × ESI method interaction term to account for the possibility that some ESI methods might perform better in specific ROIs. We computed pairwise post hoc comparisons between ESI methods for each ROI, applying Benjamini-Hochberg (Benjamini and Hochberg 1995) false discovery rate correction to control for multiple comparisons across the 37 ROIs. Results were visualized on cortical surface meshes.

## 3. Results

Throughout this section, unless stated otherwise, the results are reported in the broadband frequency band. When details are not reported in the main text, results for the remaining frequency bands are provided in the Supporting Information.

### 3.1 Dataset

The dataset included data from 9 patients collected by the Epilepsy Unit of University Hospitals of Geneva, Switzerland, and from 25 patients provided by the Niguarda Hospital of Milan, Italy. However, two patients from the first center and three patients from the second center were excluded due to bad EEG quality, leaving a final dataset of 29 patients. The median duration of the simultaneous recordings was 13.55 minutes, with an interquartile range (IQR) of 6.71 minutes. The median number of electrodes placed in gray matter, which were used in our analysis, was 79 per patient, with an IQR of 34.

Figure 2A shows the electrodes in gray matter across all patients after co-registration onto the Montreal Neurological Institute (MNI) template, in sagittal, coronal, and axial planes. This visualization illustrates the spatial coverage and reveals that, while most brain areas are sampled, certain cortical regions remain undersampled, including the superior frontal gyrus, the sensorimotor cortex, the superior parietal lobule, and the occipital lobe. Figure 2B–C show that the temporal lobe and hippocampus are more extensively sampled, reflecting the predominance of temporal lobe epilepsy in the patient cohort, which is consistent with the known prevalence of this epilepsy type.

**Figure 2.**
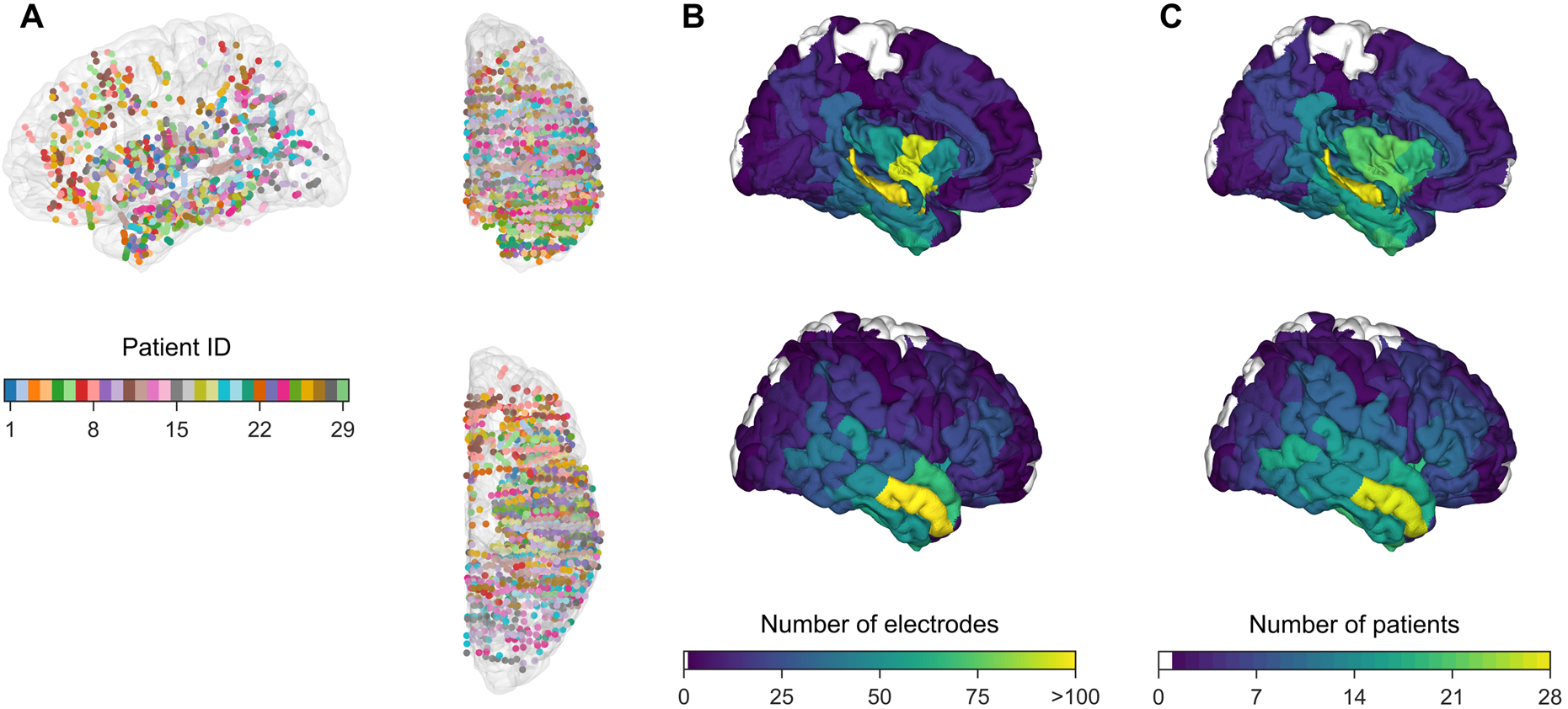
Spatial distribution of iEEG electrodes and regional coverage across patients. (A) Brain sampling across patients, illustrated by the distribution of intracranial depth electrodes located in gray matter and projected onto the MNI template brain, shown in sagittal, coronal, and axial views. All electrodes were projected onto the right hemisphere for visualization. (B) Number of electrodes per region of interest (ROI) of the Lausanne 2018 scale-3 atlas. (C) Number of patients contributing at least one electrode to each ROI. In (B) and (C), medial (top) and lateral (bottom) views are shown, and ROIs containing no electrodes are shown in white.

### 3.2 Effects of frequency band and ESI method on source reconstruction accuracy

#### 3.2.1 Model fit and interaction

We computed the iEEG–virtual iEEG correlation for each channel and patient (Figure 3A1). We modeled these values using the GLME model described in Section 2.5.1, which showed a good fit to the data (R² = 0.44). ANOVA revealed significant main effects of frequency band and ESI method, and a significant interaction between the two (all *p* < 0.001), indicating that the effect of the ESI method on correlation differed across frequency bands. Overall, estimated iEEG–virtual iEEG correlations were low across methods and frequency bands, indicating limited reconstruction of individual intracranial time series. The highest estimated correlation was 0.126, for sLORETA in the delta band. At the single-channel level, and taking eLORETA for broadband as an example, only a median of 6.4% of channels per patient (IQR 0.8–11.3%) had an observed correlation above 0.3. Estimated correlations for all frequency × method combinations are presented in Table S2.

**Figure 3.**
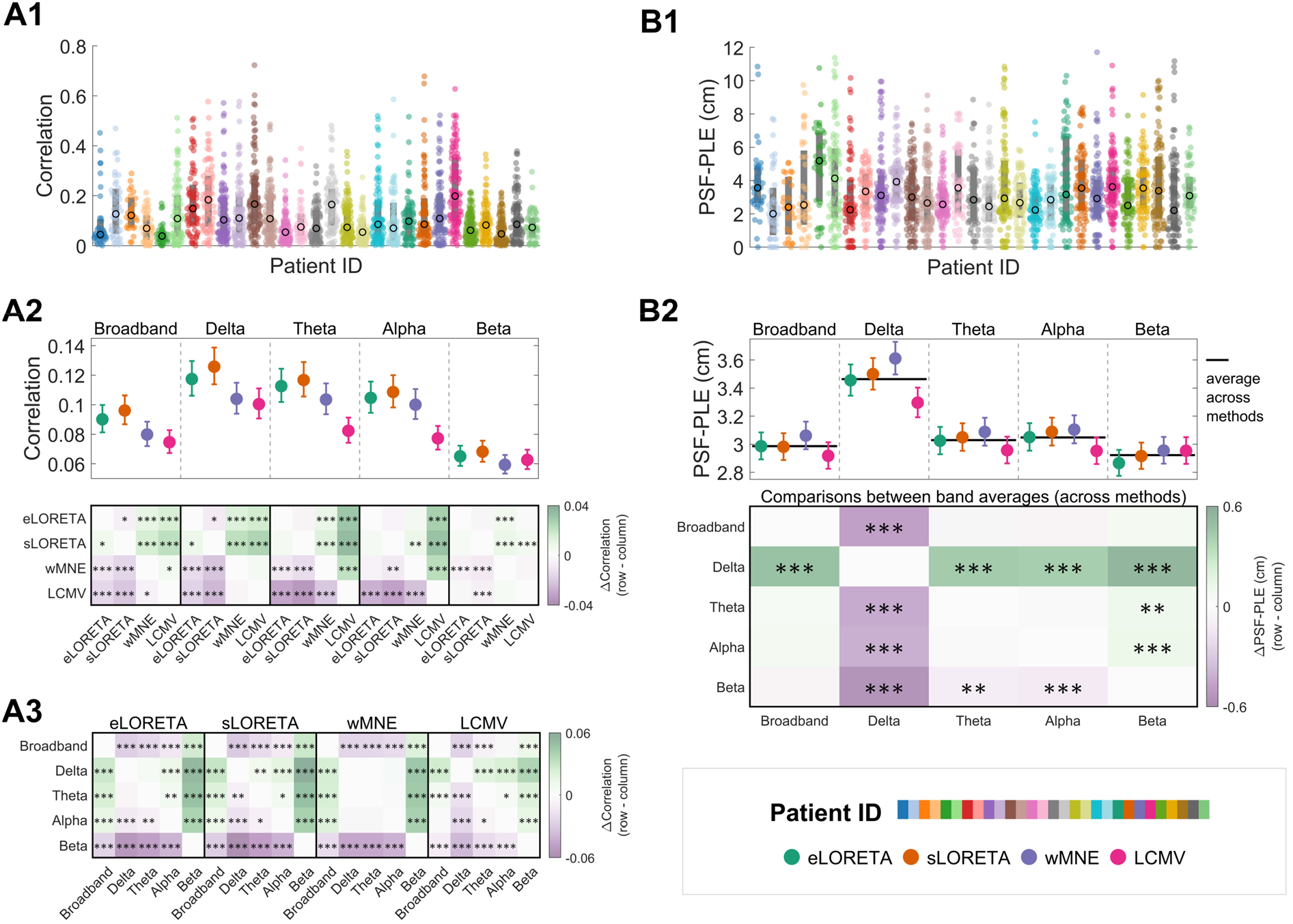
Distributions, estimated marginal means and pairwise comparisons of correlation and PSF-PLE across ESI methods and frequency bands. (A1, B1) Absolute iEEG-virtual iEEG correlation and PSF-PLE across channels (dots) and patients (colors), shown for eLORETA in the broadband range as an example. Black circles indicate the median per patient. (A2) Estimated marginal means from generalized linear mixed-effects models (GLME), back-transformed to the response scale, shown for each ESI method within each frequency band (colors denote ESI methods; error bars indicate 95% confidence intervals). The matrix below shows all pairwise comparisons between ESI methods within each frequency band, one block per band, aligned with the estimates above. (A3) Pairwise comparisons between frequency bands within each ESI method (one block per method), computed on the same estimates as in A2. (B2) Estimated marginal means for PSF-PLE, shown as in A2. As the frequency × method interaction was not significant, black horizontal lines indicate the frequency band average across ESI methods, and the single matrix below compares these band averages. In every matrix, cell color shows the difference between the values of its row and its column; color scales differ between panels. Asterisks indicate significant post hoc pairwise comparisons (*, *p* < 0.05; **, *p* < 0.01; ***, *p* < 0.001). Abbreviations as in Figure 1.

#### 3.2.2 Frequency comparisons within ESI methods

For eLORETA, sLORETA, and LCMV, a consistent ordering emerged, with delta performing best, followed by theta, then alpha, then broadband, and finally beta showing the lowest correlation in all cases. wMNE showed a somewhat flatter pattern, with delta, theta, and alpha performing comparably, all outperforming broadband and beta. Broadband outperformed beta across all methods. Detailed pairwise comparisons of frequency bands within each ESI method are presented in Table S3 and in Figure 3A3.

#### 3.2.3 ESI method comparisons within frequency bands

Across all frequency bands, sLORETA and eLORETA achieved the highest correlation, with sLORETA generally ranking first. LCMV showed the weakest performance in most bands, particularly for broadband, delta, theta, and alpha, while wMNE occupied an intermediate position. The exception was the beta band, where method differences were minimal and LCMV performed comparably to eLORETA. Despite these differences between ESI methods, the effects were small, with the largest between- method difference being only 0.034 (sLORETA vs. LCMV, theta band). All pairwise comparisons of ESI methods within each frequency band are reported in Table S4 and in Figure 3A2.

### 3.3 Effects of frequency band on the reconstruction metrics

Taking again eLORETA for broadband as an example, a median of 3.9% of channels per patient (IQR 2.6– 5.2%) had a peak localization error of the point-spread functions (PSF-PLE) of 0 cm, i.e., the maximum correlation occurred at the corresponding virtual electrode. For PSF-PLE, an ANOVA of the fitted GLME (R² = 0.41) revealed a non-significant interaction between frequency band and ESI method. The main effect of frequency band was significant (*p* < 0.001), while the main effect of ESI method was non-significant (*p* = 0.096). Post hoc comparisons averaged across methods revealed that delta showed the highest PLE, significantly greater than broadband, theta, alpha, and beta (differences of 0.478, 0.435, 0.416, and 0.541 cm, respectively, all *p* < 0.001). Both theta and alpha showed significantly higher PLE than beta (differences of 0.107 and 0.126 cm, respectively, *p* < 0.01). Overall, beta and broadband showed the lowest PSF-PLE and did not differ significantly from each other, followed by theta and alpha, with delta showing the highest PSF-PLE. Estimated values per frequency band are reported in Table S5 and detailed pairwise results are in Table S6 and in Figure 3B2.

While the effect of frequency band on PSF-PLE was consistent across ESI methods, the remaining metrics—the spatial deviation of the point-spread functions (PSF-SD) and the peak localization error and spatial deviation of the cross-talk functions (CTF-PLE, CTF-SD)—showed significant interactions between ESI method and frequency band (all *p* < 0.001; R² = 0.50, 0.41, 0.54, respectively). Post hoc comparisons were therefore conducted both across frequency bands within each ESI method and across ESI methods within each frequency band.

For PSF-SD, beta consistently showed the lowest PSF-SD for eLORETA and sLORETA, with broadband intermediate between beta and the lower bands (delta, theta, alpha). wMNE again showed the least spread in beta, but the greatest in delta, with broadband, theta, and alpha falling in between. LCMV showed a distinct pattern, with broadband exhibiting the least spread, while the remaining bands did not differ significantly (Table S7, Figure S1-A2).

For CTF-PLE, again taking eLORETA for broadband as an example, a median of 5.6% of channels per patient (IQR 4.0–8.1%) had a CTF-PLE of 0 cm, i.e., the virtual electrode was most strongly driven by its corresponding iEEG electrode. Delta consistently showed the highest values for eLORETA, sLORETA, and wMNE, with beta and broadband exhibiting the lowest values. Alpha and theta fell in an intermediate position, with alpha higher than theta. LCMV deviated from this pattern: broadband and theta performed best, while the remaining bands did not differ significantly from one another (Table S7, Figure S1-B2).

For CTF-SD, theta and broadband consistently exhibited the lowest values across methods, while delta and beta showed the highest. eLORETA and sLORETA yielded nearly identical orderings, with theta performing best and delta and beta performing worst. wMNE followed a similar trend, though delta, alpha, and beta were more comparable with one another. LCMV behaved differently: beta emerged as the worst-performing band by a large margin, while theta and broadband again performed best (Table S7, Figure S1-C2).

### 3.4 Effects of ESI method on the reconstruction metrics

#### 3.4.1 Spatial deviation of the point-spread functions

For spatial deviation of the point-spread functions (PSF-SD), across all frequency bands, eLORETA consistently exhibited the lowest values. sLORETA generally followed as the second-best method, performing comparably to eLORETA for theta and alpha but showing significantly higher PSF-SD for the remaining bands. The relative performance of wMNE and LCMV varied by band: wMNE showed the highest PSF-SD for broadband, delta, and theta, while LCMV showed the highest PSF-SD for beta. For alpha, wMNE and LCMV performed similarly (Table S8, Figure S1-A3).

#### 3.4.2 Peak localization error of the cross-talk functions

For peak localization error of the cross-talk functions (CTF-PLE), across frequency bands, no consistent method ranking emerged. LCMV performed best for delta and theta but worst for beta, while wMNE showed the greatest localization error for theta and alpha. eLORETA performed consistently well, never showing the highest CTF-PLE in any band. For broadband, all methods performed comparably (Table S8, Figure S1-B3).

#### 3.4.3 Spatial deviation of the cross-talk functions

For spatial deviation of the cross-talk functions (CTF-SD), across frequency bands, eLORETA and sLORETA consistently exhibited the lowest values, outperforming wMNE for theta and alpha, and all other methods for broadband. For delta, all methods performed comparably. Beta showed the most differentiation, with eLORETA performing best and LCMV worst, while sLORETA and wMNE fell in between (Table S8, Figure S1-C3).

Estimated values of the reconstruction metrics derived from the three GLME models for all combinations of frequency band and ESI method are presented in Table S9.

#### 3.4.4 ESI method ranking

Using the Copeland-style scoring system, the ESI methods were ranked for each frequency band as follows: for PSF-based metrics, eLORETA ranked first in the delta, theta, and beta bands and tied with sLORETA in broadband, while sLORETA ranked first in the alpha band, followed by eLORETA. The third position was occupied by wMNE in the alpha and beta bands, while wMNE and LCMV tied in broadband. For CTF-based metrics, eLORETA ranked first in the theta, alpha, and beta bands, while sLORETA ranked first in the broadband and delta bands. The third position was occupied by LCMV in the delta, theta, and alpha bands and by wMNE in the broadband and beta bands. Detailed results are presented in Table S10.

### 3.5 Relationship between reconstruction metrics and iEEG-virtual iEEG correlation

Using the GLME models described in Section 2.5.2, we assessed the relationship between the reconstruction metrics and the iEEG-virtual iEEG correlation. Results were largely consistent across methods; for clarity, we describe the broadband results for eLORETA in detail, with complete fixed- effect estimates for all methods and frequency bands reported in Figure 4 and Table S11.

**Figure 4.**
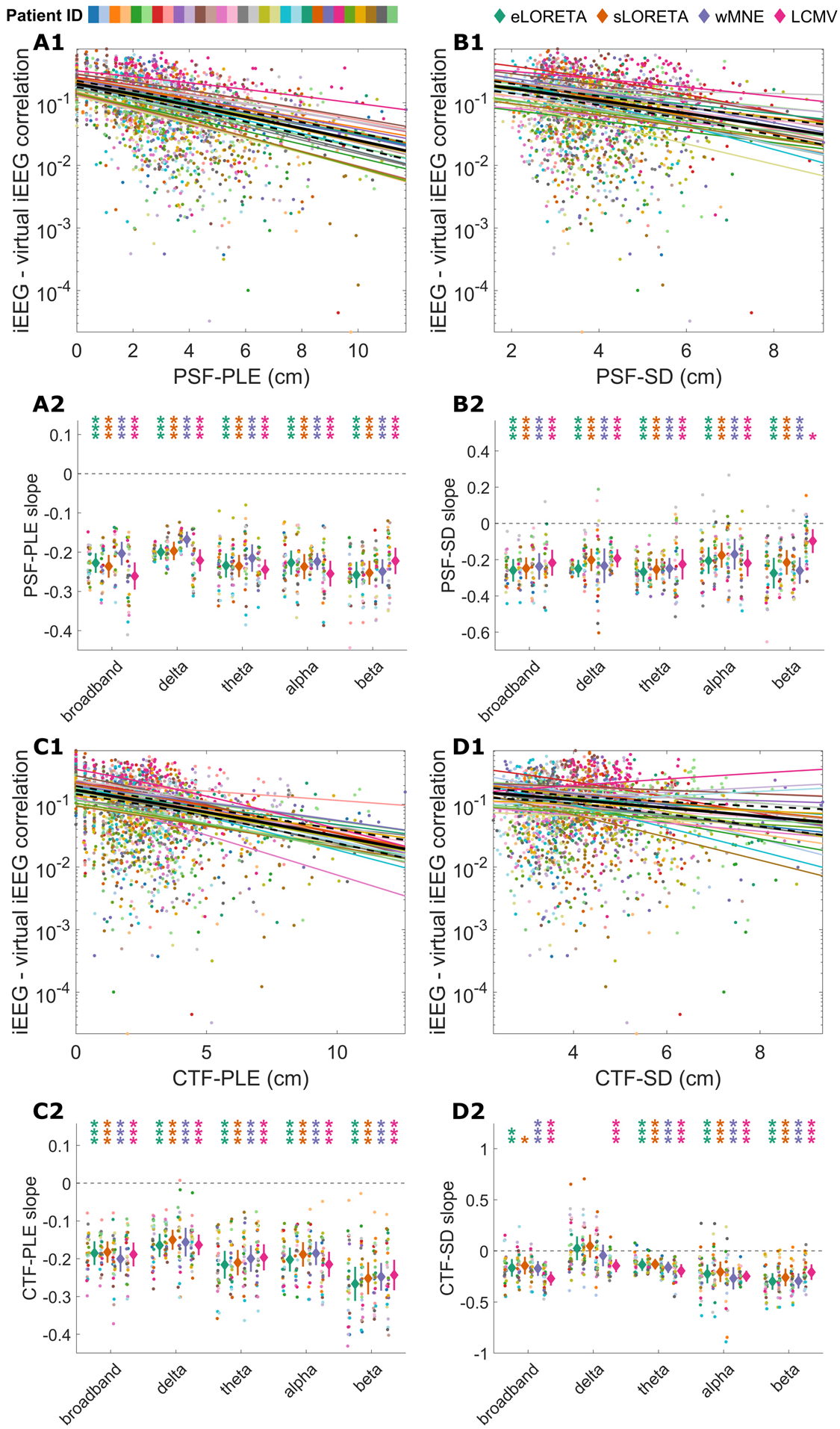
Relationship between iEEG- virtual iEEG correlation and PSF- and CTF-based reconstruction metrics. (A1, B1) Scatter plots of absolute correlation values (log scale) versus PSF-PLE and PSF-SD, shown for eLORETA in the broadband range as an example; each dot represents one iEEG-virtual iEEG electrode pair, with colors indicating patient identity. Colored lines show patient-specific slopes of the generalized linear mixed-effects (GLME) models. Black solid and dashed lines indicate the population-level fixed effects and their 95% confidence intervals. (A2, B2) Fixed-effect slopes (diamonds with lines indicating the 95% confidence intervals) and the corresponding patient-specific random slopes (colored points) for the PSF-PLE and PSF-SD across ESI methods and frequency bands. (C1, D1, C2, D2) Same as (A1, B1, A2, B2), but for CTF-PLE and CTF-SD. Asterisks indicate fixed-effect slopes significantly different from zero, colored to match the corresponding ESI method (*, *p* < 0.05; **, *p* < 0.01; ***, *p* < 0.001). Abbreviations as in Figure 1.

For PSF metrics, both PLE and SD showed significant negative fixed-effect slopes across all ESI methods. For eLORETA, PSF-PLE yielded β = −0.228 (95% CI: [−0.253, −0.202], p < 0.001, R² = 0.415), and PSF-SD yielded β = −0.259 (95% CI: [−0.321, −0.196], p < 0.001, R² = 0.217). These negative relationships were largely consistent across all five frequency bands for both metrics, indicating that greater localization error and leakage were associated with poorer reconstruction quality (Figure 4A2, B2).

For CTF metrics, similar patterns were observed. For eLORETA, CTF-PLE yielded β = −0.185 (95% CI: [−0.217, −0.154], *p* < 0.001, R² = 0.297), and CTF-SD yielded β = −0.166 (95% CI: [−0.252, −0.080], *p* < 0.01, R² = 0.228). These negative relationships were largely consistent across frequency bands and methods, except for CTF-SD in the delta band for eLORETA, sLORETA, and wMNE, for which the effect was not significant (Figure 4C2, D2).

### 3.6 Electrode depth and spike rate

Using GLME models, we assessed the effect of electrode depth and spike rate on the iEEG-virtual iEEG correlation. Results across all methods and frequency bands are shown in Figure 5 and Table S12. For electrode depth, a significant negative fixed-effect slope was observed in the broadband signal for eLORETA (β = −0.175, 95% CI: [−0.245, −0.106], *p* < 0.001, R² = 0.247), indicating that electrodes located further from the scalp surface were associated with poorer reconstruction quality. This negative effect was consistent across all ESI methods and frequency bands (Figure 5A).

**Figure 5.**
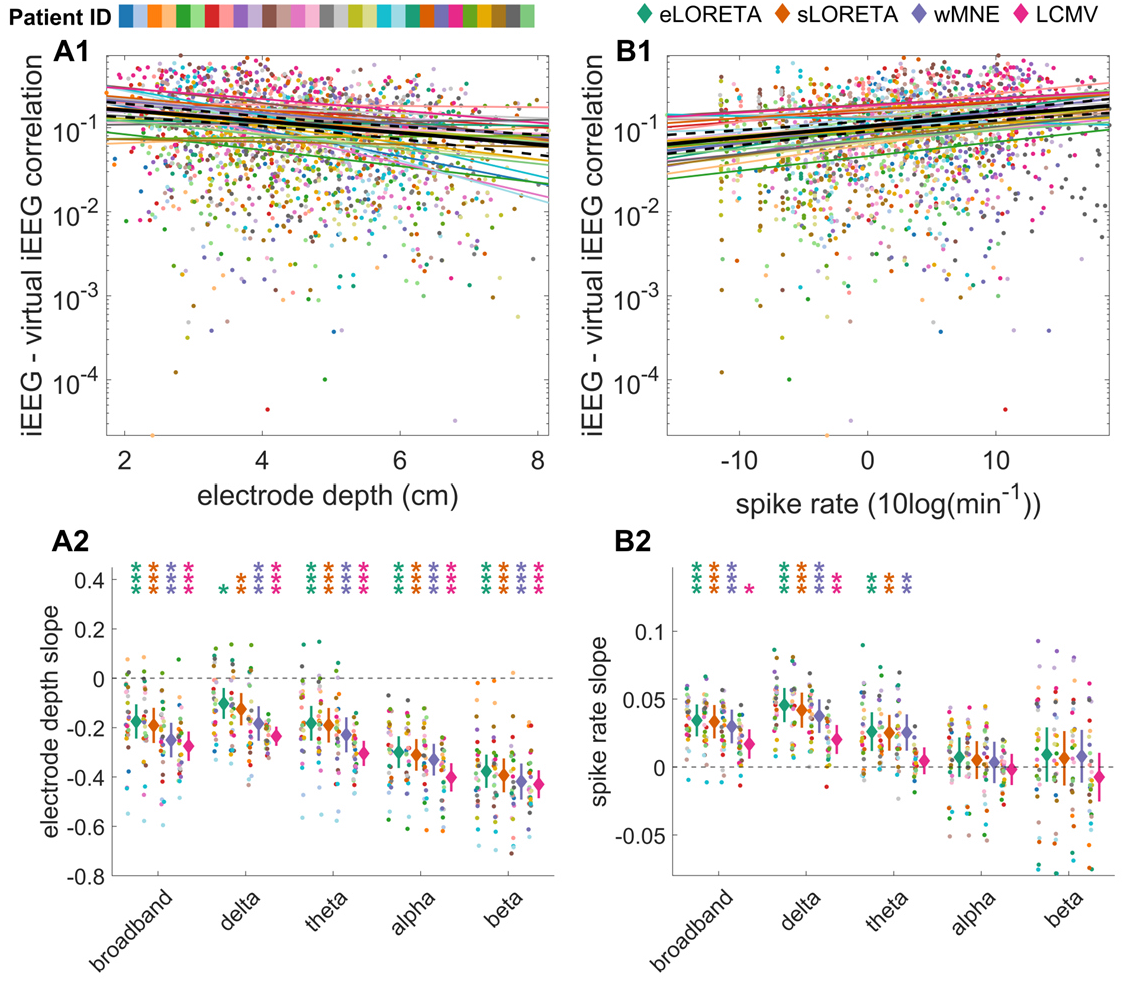
Relationship between intracranial data properties (electrode depth and spike rate) and iEEG-virtual iEEG correlation. In panels A1 and B1, each dot represents one iEEG-virtual iEEG electrode pair, and colors indicate patient identity. (A1) Scatter plot of absolute correlation values (log scale) versus electrode depth, shown for eLORETA in the broadband range as an example. Colored lines show patient-specific slopes of the generalized linear mixed-effects (GLME) models. Black solid and dashed lines indicate the population-level fixed effect and its 95% confidence interval. (A2) Fixed-effect slopes (diamonds colored by ESI method, with lines indicating the 95% confidence intervals) and the corresponding patient-specific random slopes (colored points) for the electrode depth model across ESI methods and frequency bands. (B1, B2) Same as (A1, A2), but for spike rate (in 10 log4o(spikes/min)). Asterisks indicate fixed-effect slopes significantly different from zero, colored to match the corresponding ESI method (*, *p* < 0.05; **, *p* < 0.01; ***, *p* < 0.001). Abbreviations as in Figure 1.

For spike rate, a significant positive fixed-effect slope was observed for eLORETA (β = 0.034, 95% CI: [0.023, 0.046], *p* < 0.001, R² = 0.190), indicating that a higher spike rate was associated with better reconstruction quality. The positive effect was also consistent in the delta and theta bands (Figure 5B).

### 3.7 Performance across regions of interest

GLME models for the broadband showed a good fit across all five metrics (R² ranging from 0.585 to 0.851). Significant ROI × ESI method interactions were observed across all metrics and frequency bands (all *p* < 0.001). Full results are reported in Table S13. A direct comparison between eLORETA and sLORETA showed significant differences in only four ROIs (Table S14). We therefore present the comparisons of each of them against wMNE and LCMV separately.

For correlation, eLORETA generally performed better than LCMV, particularly in temporal, insular and medial temporal regions (including the hippocampus, parahippocampal gyrus, entorhinal cortex and amygdala). In the theta and alpha bands, these differences extended more posteriorly. In a few frontal and central regions, LCMV showed higher correlation. In the beta band, differences were more limited.

The PLE results were less consistent. For PSF-PLE, eLORETA showed higher PSF-PLE than LCMV in several lateral temporal regions, but lower PSF-PLE than wMNE in the hippocampus in most frequency bands. For CTF-PLE, a similar distinction appeared between lateral and medial regions: eLORETA tended to show higher error than LCMV in some small areas, such as the rostral middle frontal region, but lower error in deeper structures, such as the hippocampus.

The SD results were more consistent. Both PSF-SD and CTF-SD showed lower SD values for eLORETA across many regions, indicating more focal source estimates. This effect was most evident in the broadband, delta and beta bands for PSF-SD and in the beta band for CTF-SD and involved a wide range of cortical and medial temporal areas, as shown in Figure 6. In the other frequency bands, the effect was still present but more limited.

**Figure 6.**
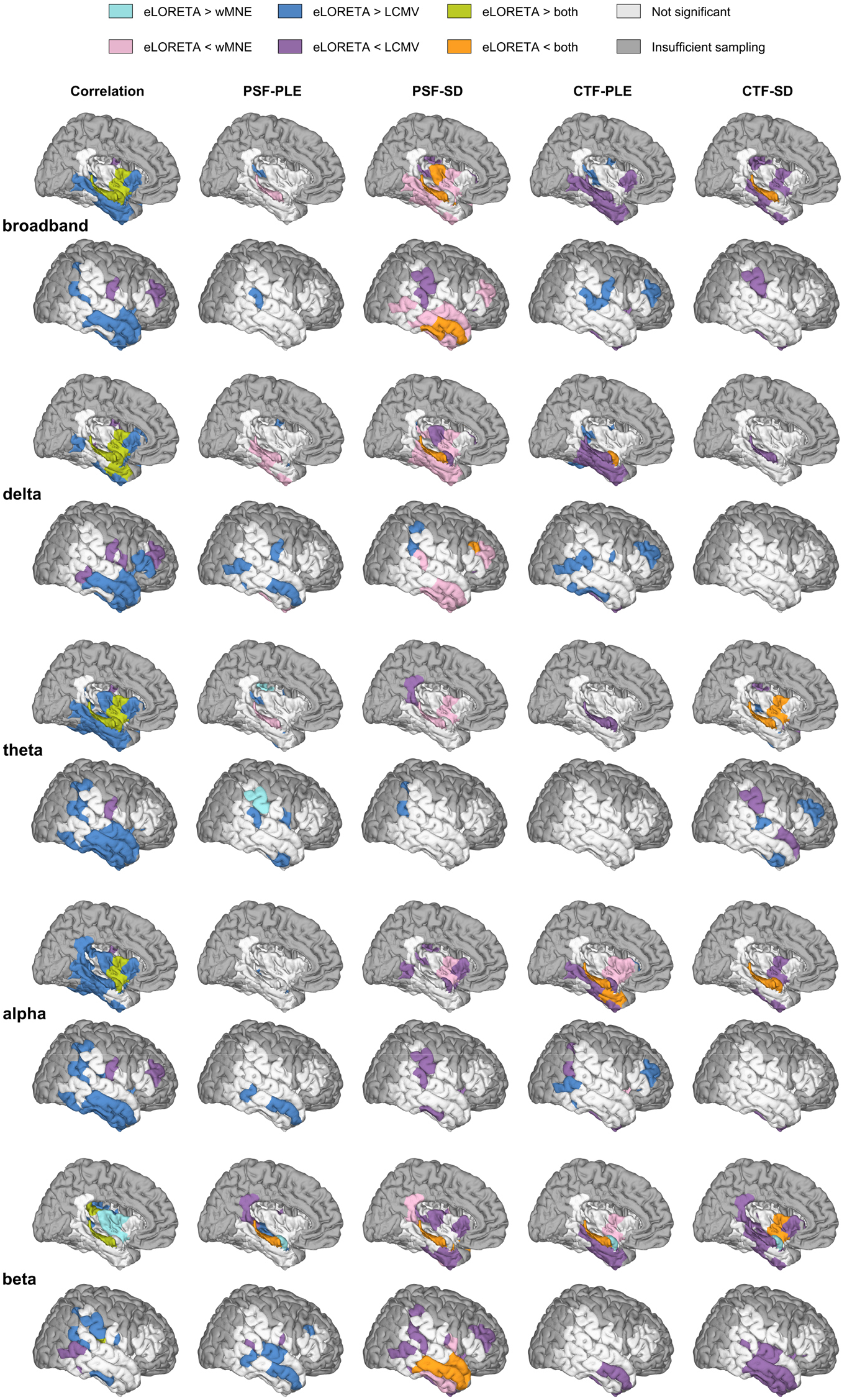
ROI-wise comparison of eLORETA against wMNE and LCMV across frequency bands, for correlation and reconstruction metrics. Right-hemisphere medial (upper) and lateral (lower) views of the Lausanne 2018 scale-3 atlas, shown for the broadband, delta, theta, alpha, and beta frequency bands (rows, top to bottom). Columns show the iEEG-virtual iEEG correlation and the four reconstruction metrics (PSF-PLE, PSF-SD, CTF-PLE, and CTF- SD). Higher correlation but lower PLE/SD reflects better reconstruction. For each ROI, pairwise comparisons between eLORETA and the two reference methods (wMNE, LCMV) were performed on the estimated values; *p*- values were FDR-corrected across ROIs (Benjamini–Hochberg, *q* < 0.05). ROIs with insufficient patient sampling (< 8 patients) are shown in dark gray. Abbreviations as in Figure 1.

Overall, method differences varied across ROIs but did not appear to be confined to specific lobes or regions, suggesting that the observed differences were not driven exclusively by anatomical features.

An equivalent analysis comparing sLORETA with wMNE and LCMV yielded similar patterns, with method differences more pronounced for correlation than for spatial accuracy metrics (Figure S2).

## 4. Discussion

Using simultaneous hdEEG-iEEG recordings, we demonstrated that ESI can recover aspects of cortical spontaneous activity, but the fidelity of reconstruction remains limited when evaluated at the level of individual time series. We also presented a comprehensive characterization of four major ESI methods in terms of their relationship to physiological factors and canonical reconstruction metrics.

### 4.1 Source-reconstructed EEG only partially captures local spontaneous cortical dynamics

Although some channels showed higher correlations (up to about 0.9, for sLORETA in the alpha band), the overall correlation between virtual and actual iEEG signals was low, with an estimated correlation of 0.126 in the best-performing case (sLORETA, delta band). For eLORETA in broadband, only a median of 6.4% of channels per patient exceeded a correlation of 0.3. Importantly, this metric is more stringent than simply assessing signal similarity, as it also evaluates whether the intracranial time series is reconstructed at the correct anatomical location (diagonal of the correlation matrix). Consequently, low correlations may arise not only from inaccurate reconstruction of the underlying neural dynamics but also from mislocalization of the reconstructed activity. Consistent with this interpretation, localization errors were approximately 3 cm across most frequency bands, and only a median of 3.9% of channels per patient had a zero PLE (eLORETA, broadband), indicating that the reconstructed signal usually showed greater similarity to the intracranial recording at locations other than that of the corresponding electrode. These findings suggest that improving the spatial accuracy of source reconstruction is likely to enhance the reconstruction of spontaneous cortical time series.

An important distinction between the present study and previous ESI validation studies is that we evaluated the reconstruction of neural time series rather than localization accuracy alone (van Mierlo et al. 2017; Coito et al. 2019; Sohrabpour et al. 2020; Hauk et al. 2022; Pascarella et al. 2023; Unnwongse et al. 2023; Zauli et al. 2024). These studies typically focused on focal and time-constrained events, such as epileptic spikes (van Mierlo et al. 2017; Coito et al. 2019; Sohrabpour et al. 2020; Zauli et al. 2024) or electrical stimulation (Pascarella et al. 2023; Unnwongse et al. 2023), which generally provide higher-SNR signals and can often be averaged to further improve the SNR. In contrast, the present study evaluated the reconstruction of spontaneous ongoing activity arising from multiple simultaneously active brain regions, a substantially more challenging task in which averaging cannot be used to improve the SNR.

A further difficulty is that, unlike time-locked paradigms with a defined pre-stimulus baseline period, spontaneous recordings do not provide an obvious baseline period for estimating the noise covariance used in some ESI approaches. To our knowledge, this is the first study to comprehensively evaluate the ability of commonly used ESI methods to reconstruct spontaneous cortical time series using simultaneous hdEEG-iEEG recordings. Rather than questioning the established utility of ESI for localizing focal or evoked activity, our findings indicate that accurately reconstructing the temporal dynamics of spontaneous neural activity at individual locations remains considerably more challenging. This distinction is particularly important for applications that rely on source-reconstructed time series, such as functional connectivity analyses, where limited temporal fidelity and spatial leakage may influence the interpretation of the results. This may partly explain why a recent study comparing MEG source imaging with simultaneous iEEG found only moderate to low agreement between MEG- and iEEG-derived functional connectomes (Afnan et al. 2026).

### 4.2 Physiological factors influence reconstruction fidelity

In our study, reconstruction quality, as measured by the iEEG-virtual iEEG correlation, decreased with increasing iEEG electrode depth. This effect was observed across all ESI methods and frequency bands, suggesting a general limitation in reconstructing deep compared to superficial brain activity. This finding is expected, given the reduced sensitivity of scalp EEG to deep sources, which generate weaker and spatially broader scalp voltage maps. A similar relationship between electrode depth and spatial accuracy has been reported in both simulation and stimulation-based studies (Mikulan et al. 2020; Unnwongse et al. 2023). We also observed no consistent ROI-specific advantage for any ESI method across the analyzed ROIs, suggesting that reconstruction performance may be driven more by factors such as source depth and individual anatomical geometry than by the anatomical location itself.

Moreover, reconstruction quality increased with spike rate, likely reflecting higher signal power, as epileptic spikes arise from the synchronous activity of spatially extended neuronal populations and are thus associated with larger EEG amplitudes (Uhlhaas and Singer 2006; Cohen 2017). Although spike rate was examined here because of the pathology of the studied patients, we hypothesize that the same principle may extend to other transient graphoelements, such as bursts of oscillatory activity, because they similarly arise from synchronous neuronal populations. In contrast, the ongoing aperiodic background activity underlying the 1/f component is thought to reflect the average activity of asynchronous local neuronal populations, resulting in weaker scalp signals due to partial cancellation through volume conduction (Miller et al. 2009). These already attenuated signals are further smoothed by the ESI operator, making them more difficult to reconstruct accurately. Consequently, recordings with a higher proportion of transient events relative to background activity are expected to yield better reconstruction performance. Future studies comparing reconstruction performance during interictal and ictal periods could further test this hypothesis, as seizures are generally characterized by more sustained, high-amplitude, and synchronized neuronal activity than the interictal period.

Reconstruction performance tended to decrease with increasing frequency, with lower-frequency activity (delta and theta) being better reconstructed than higher-frequency activity. This pattern may partly reflect differences in signal amplitude, as the power of neural signals follows an approximate 1/f distribution, resulting in greater power at lower frequencies. In addition, lower-frequency activity may be more easily reconstructed because it often reflects larger-scale synchronized neuronal activity, which generates stronger and more spatially coherent scalp signals (Buzsáki 2006). In our data, broadband signals combine multiple frequency components with distinct amplitudes and spatial characteristics. After source reconstruction, each frequency component may undergo different degrees of spatial leakage, and the resulting mixture may therefore reduce broadband reconstruction accuracy compared with narrow frequency bands such as delta and theta. Consistent with this observation, previous work combining an iEEG atlas and MEG recording of healthy participants showed that removing the aperiodic component can improve the power spectrum concordance of the two modalities (Afnan et al. 2023). This finding highlights that broadband source-reconstructed signals should be interpreted more cautiously, particularly for analyses relying on accurate characterization of spectral properties, including aperiodic activity or relative spectral power.

### 4.3 Spatial resolution constrains time-series reconstruction

PSF- and CTF-based metrics provide complementary information and should both be considered when characterizing source reconstruction performance. In the context of this study, PSFs describe how much the activity recorded by an actual iEEG electrode spreads to virtual electrodes, whereas CTFs reflect how much the signal at a given virtual iEEG electrode is influenced by activity from actual iEEG electrodes.

For both PSF- and CTF-based peak localization error (PLE) and spatial deviation (SD), we observed a negative relationship with iEEG-virtual iEEG correlation across all ESI methods and frequency bands, except for CTF-SD in the delta band for eLORETA, sLORETA, and wMNE. This suggests that poorer spatial accuracy and increased leakage were associated with less accurate reconstruction of the time series.

At first glance, the frequency-dependent results, namely the fact that the frequency band with the lowest PLE/SD did not necessarily show the highest iEEG-virtual iEEG correlation, appear inconsistent with the above negative relationship. However, these findings address different questions. The negative relationship was observed within each frequency band, where electrodes with lower PLE or SD tended to show higher iEEG-virtual iEEG correlation. In contrast, comparisons between frequency bands are also influenced by the intrinsic spatial characteristics of the underlying neural activity. For example, PSF-based metrics favored the beta band despite its poor time-series reconstruction, possibly because beta activity arises from more spatially localized neuronal populations (Buzsáki 2006), producing PSFs with a few high values surrounded by a relatively flat profile. Such distributions naturally yield lower SD and can also reduce PLE, independently of how accurately the underlying time series is reconstructed. Consequently, it is not surprising that the frequency band with the best spatial resolution metrics was not necessarily the one with the highest time-series reconstruction accuracy.

Overall, these results suggest that leakage is not merely a spatial limitation of ESI but also a determinant of the fidelity of reconstructed neural dynamics. Consequently, methods or parameter settings that reduce leakage are likely to be important for applications relying on source-reconstructed time series. This interpretation is consistent with previous studies showing that post hoc leakage correction improves the recovery of underlying functional connectivity (Colclough et al. 2015; Farahibozorg et al. 2018).

### 4.4 No universally optimal ESI approach

Across most frequency bands, eLORETA achieved the best overall performance when combining time- series reconstruction and spatial accuracy metrics (Copeland score). However, examining each metric separately revealed a trade-off between reconstruction properties: sLORETA generally yielded higher iEEG-virtual iEEG correlations, suggesting better recovery of the signal at the corresponding electrode location, whereas eLORETA consistently showed lower spatial spread for both PSF-SD and CTF-SD. In contrast, localization error did not differ consistently between methods. These findings indicate that no single ESI method provides optimal performance across all aspects of source reconstruction, with sLORETA favoring time-series correspondence and eLORETA providing more spatially focal estimates. Overall, eLORETA may represent a balanced solution when both temporal and spatial properties are considered, although the differences between methods remain relatively modest in the context of spontaneous whole-brain activity.

Interestingly, LCMV showed the poorest time-series reconstruction across frequency bands. This may reflect limitations of beamforming in the context of spontaneous activity, where simultaneously active and correlated sources violate the assumption of uncorrelated source activity. Future studies should evaluate the effect of different corrections for the center-of-head bias and covariance estimation.

From a theoretical perspective, ESI methods based on reweighting the classical MNE solution (e.g., wMNE, sLORETA, and eLORETA) are designed to modify the PSFs of the resolution matrix but do not directly modify its CTFs (Hauk et al. 2022). This framework is consistent with the more pronounced differences observed between methods for PSF-based metrics than for CTF-based metrics. In contrast, beamforming approaches construct data-driven spatial filters, which influence both the PSFs and the CTFs. Nevertheless, modest differences in CTF-based metrics were still observed in the present study. This likely reflects the experimental setting: spontaneous, distributed whole-brain activity rather than idealized single-source conditions.

### 4.5 Methodological considerations and future directions

One methodological consideration is that skull defects due to burr holes created during depth electrode implantation were not incorporated into the head model. Although such openings may affect current propagation, finite element simulations suggest that their impact is limited (Wong et al. 2024), and previous work has shown that the number of skull openings does not substantially affect dipole fitting accuracy (Unnwongse et al. 2023). Because this modeling choice was consistent across all ESI methods, it is unlikely to have affected the relative comparison between methods.

We also used a single regularization parameter value across methods. Since regularization was matched across methods, we believe that the differences found here are unlikely to be driven solely by regularization but rather reflect intrinsic differences between the ESI operators. Future studies should therefore investigate whether optimizing regularization separately for each ESI method improves time- series reconstruction and spatial accuracy.

Finally, further studies could replicate this work using simultaneous MEG-iEEG recordings. As MEG is less affected by skull conductivity than EEG, localization accuracy—and consequently time-series reconstruction—could be improved.

These findings do not argue against the utility of ESI but rather define the spatial and temporal scales at which EEG source reconstruction should be interpreted. More broadly, they highlight the need to improve existing ESI methods or develop new ones to better reconstruct neural time series from EEG.

### Data availability statement

The raw data that support the findings of this study are available on reasonable request from the corresponding author for the Geneva dataset, and from the investigators at Niguarda Hospital, Milan, for the Milan dataset. The raw data are not publicly available as they could contain information that could compromise the privacy of research participants. The pseudonymized processed data used for the statistical analyses (per-contact correlation and reconstruction metrics, regions of interest, electrode depth and spike rate) will be made publicly available on figshare upon acceptance of the manuscript.

## Funding statement

M.D. and N.R. were supported by the Swiss National Science Foundation (SNSF) Ambizione grant 209120, awarded to N.R. A.P. and E.M. were supported by the following grants: Progetto di Ricerca di Rilevante Interesse Nazionale (PRIN, P2022FMK77), Ministry of University and Research (MUR); the Italian National Recovery and Resilience Plan (NRRP), M4C2, funded by the European Union – NextGenerationEU (Project IR0000011, CUP B51E22000150006, “EBRAINS-Italy”); HORIZON-INFRA-2022-SERV (Grant No. 101147319), “EBRAINS 2.0: A Research Infrastructure to Advance Neuroscience and Brain Health”; and the European Research Council (ERC-2022-STG - 101078497 - MYFIRSTBODY; ERC-2022-SYG - 101071900 - NEMESIS). S.V. was supported by SNSF grant 236136 and SNSF Sinergia grant 209470.

## Conflict of interest disclosure

S.V. has shares in Clouds of Care. N.R. holds an international patent for Delphos (EP3422935B1). The other authors report no competing interests. We confirm that we have read the Journal’s position on issues involved in ethical publication and affirm that this report is consistent with those guidelines.

## Ethics approval statement

This study was approved by the Regional Research Ethics Committee of Geneva (protocol 13-004) and by the Ethics Committee of Niguarda Hospital, Milan, Italy (protocol 463-092018).

## Patient consent statement

All patients provided informed consent before participating.

