## Supporting Information for "How well does electrical source imaging reconstruct neural time series? Evidence from simultaneous scalp and intracranial EEG recordings"

### Supporting Methods

Detailed formulations of the ESI methods used in this study are provided below.

Given a precomputed leadfield $L$, the minimum norm estimate (MNE) solution is given by:

$$G_{MNE} = RL^T(LRL^T + \lambda I)^{-1}$$

where $R$ is the source covariance matrix, $\lambda$ is the regularization parameter, and the noise covariance matrix is assumed to be the identity matrix as no baseline is available due to the reconstruction of spontaneous activity.

wMNE reduces the bias towards superficial sources (Lin et al. 2006) by introducing depth weights into the source covariance matrix, so that deeper sources, which have weaker leadfields, are less penalized:

$$G_{wMNE} = R_w L^T(LR_w L^T + \lambda I)^{-1}$$

where $R_w$ is a diagonal matrix with depth weights, defined here as inverse of the square of the Frobenius norm of the leadfield associated with each source.

The sLORETA (Pascual-Marqui 2002) standardizes the MNE estimates $G_{MNE}$ with the help of the normalization matrix $N_{sLORETA}$ and more specifically using the resolution matrix $R_{MNE}$:

$$G_{sLORETA} = N_{sLORETA}\ G_{MNE}$$

$$N_{sLORETA}^2 = diag(R_{MNE})^{-1}$$

where $R_{MNE}$ = $G_{MNE}\ L$ is the resolution matrix of the MNE.

Unlike wMNE, which compensates for depth bias by weighting the source covariance matrix according to leadfield strength, sLORETA normalizes each source estimate by the corresponding diagonal element of the resolution matrix, yielding theoretically zero peak localization error for point sources in the absence of noise. Unlike sLORETA, which standardizes the MNE solution, eLORETA iteratively reweights the source covariance matrix $R$ within the inverse solution itself, also yielding theoretically zero peak localization error for point sources (Pascual-Marqui 2007). To ensure fair comparisons across ESI methods, we applied the following change of variables:

$$\lambda = \alpha \frac{tr(LRL^T)}{tr(I)}$$

$tr(.)$ is the trace of a square matrix. Here, $\alpha$ is a positive coefficient that is more meaningful than $\lambda$, as it adapts to the leadfield and the prior source covariance matrix. We used the default value of $\alpha = 0.05$, as implemented in Fieldtrip. This is what we refer to as $\lambda = 5\%$ in the main text.

For the LCMV (Van Veen et al. 1997), the spatial filters were obtained using the unit-noise normalization (Westner et al. 2022) which aims to remove the center of head bias. Here, the regularization is applied during the inversion of the data covariance matrix, computed on the whole recording. As above, the regularization is written as a ratio: $C^{-1} = \left(C + \alpha \frac{tr(C)}{tr(I)} I\right)^{-1}$. We also used the default value of $\alpha = 0.05$.

### Figures

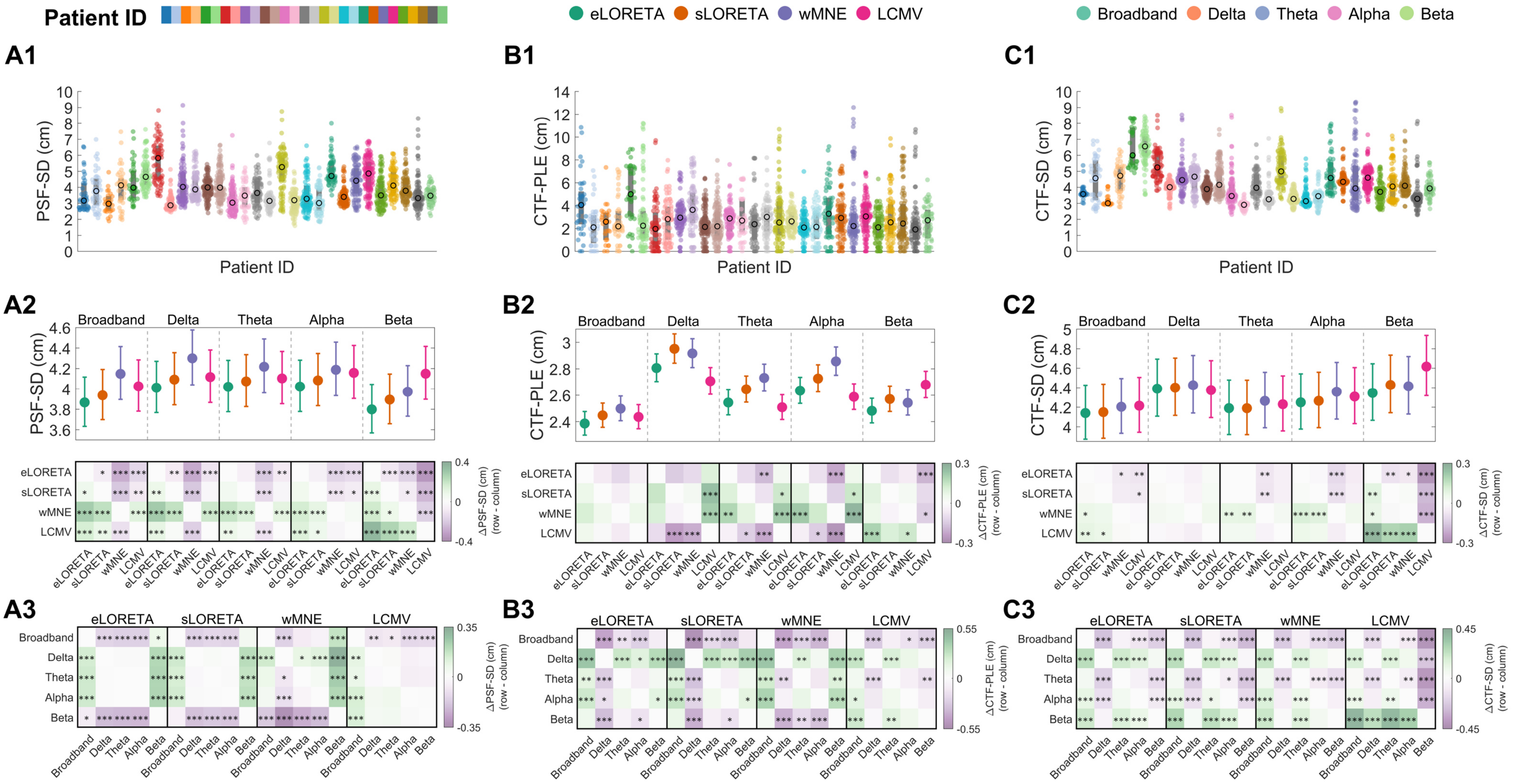


Figure S1. Distributions and pairwise comparisons of reconstruction metrics across ESI methods and frequency bands. (A1, B1, C1) Distributions of PSF-SD, CTF-PLE and CTF-SD across channels (dots) and patients (colors). Black circles indicate the median per patient. (A2, B2, C2) Estimated marginal means from generalized linear mixed-effects models (GLME), back-transformed to the response scale, for each ESI method within each frequency band (colors denote ESI methods; error bars indicate 95% confidence intervals). The matrices below show all pairwise comparisons between ESI methods within each frequency band, one block per band, below the corresponding estimates. (A3, B3, C3) Pairwise comparisons between frequency bands within each ESI method (one block per method), computed on the same estimates as in A2, B2, and C2. In every matrix, cell color shows the difference between the values of its row and its column; color scales differ between panels. Asterisks indicate significant post hoc pairwise comparisons (*, $p < 0.05$; **, $p < 0.01$; ***, $p < 0.001$). Abbreviations as in Figure 1 of the main text.

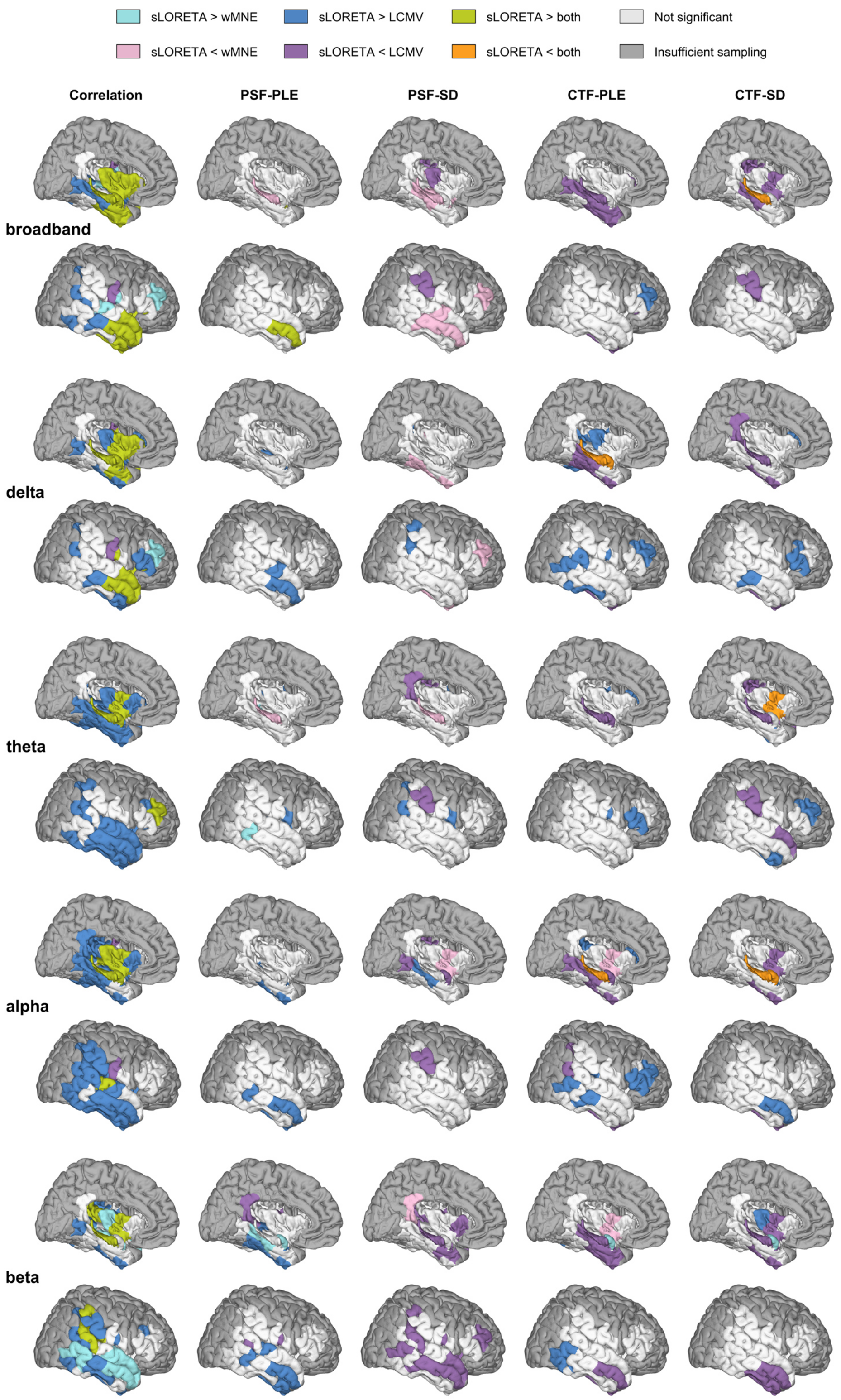


Figure S2. ROI-wise comparison of sLORETA against wMNE and LCMV across frequency bands, for correlation and reconstruction metrics. Right-hemisphere medial (upper) and lateral (lower) views of the Lausanne 2018 scale-3 atlas, shown for the broadband, delta, theta, alpha, and beta frequency bands (rows, top to bottom). Columns show the iEEG-virtual iEEG correlation and the four reconstruction metrics (PSF-PLE, PSF-SD, CTF-PLE, and CTF-SD). Higher correlation but lower PLE/SD reflects better reconstruction. For each ROI, pairwise comparisons between sLORETA and the two reference methods (wMNE, LCMV) were performed on the estimated values; *p*-values were FDR-corrected across ROIs (Benjamini-Hochberg, $q < 0.05$). ROIs with insufficient patient sampling (< 8 patients) are shown in dark gray. Abbreviations as in Figure 1 of the main text.

### Tables

| Patient | Excluded | Sex | Age at recording | Type of epilepsy | MRI findings | Electrode manufacturer |
|---|---|---|---|---|---|---|
| 1 | NO | F | 27 | LPLE | FCD | AD-Tech |
| 2 | NO | M | 48 | RTLE | None | AD-Tech |
| 3 | NO | F | 53 | RTLE | None | AD-Tech |
| 4 | YES | M | 38 | BTLE | None | AD-Tech |
| 5 | NO | F | 42 | LTLE | HS | AD-Tech |
| 6 | NO | M | 21 | BTLE | None | AD-Tech |
| 7 | NO | M | 50 | BTLE | HS | AD-Tech |
| 8 | YES | M | 27 | RTLE | None | AD-Tech |
| 9 | NO | M | 40 | LTLE | HS | Dixi Medical |
| 10 | NO | M | 21 | LFLE | None | Dixi Medical |
| 11 | NO | M | 36 | Multifocal | None | Dixi Medical |
| 12 | NO | M | 46 | RT+LE | None | Dixi Medical |
| 13 | NO | F | 30 | LFLE | None | Dixi Medical |
| 14 | NO | F | 50 | RTLE | Bilateral PNH | Dixi Medical |
| 15 | NO | M | 44 | RTLE | None | Dixi Medical |
| 16 | NO | F | 39 | LTLE | None | Dixi Medical |
| 17 | NO | M | 19 | ROLE | None | Dixi Medical |
| 18 | NO | F | 20 | LTLE | None | Dixi Medical |
| 19 | NO | F | 28 | Unknown | None | Dixi Medical |
| 20 | NO | M | 35 | LTLE | None | Dixi Medical |
| 21 | NO | F | 40 | LT+LE | None | Dixi Medical |
| 22 | YES | M | 32 | BTLE | Bilateral PNH | Dixi Medical |
| 23 | NO | F | 34 | RTLE | Right PNH | Dixi Medical |
| 24 | YES | F | 46 | LTLE | Left temporo-basal FCD | Dixi Medical |
| 25 | NO | M | 46 | BTLE | Right PNH, left hippocampal malformation | Dixi Medical |
| 26 | NO | M | 35 | RTLE | None | Dixi Medical |
| 27 | NO | F | 32 | RTLE | None | Dixi Medical |
| 28 | YES | M | 32 | RTLE | None | Dixi Medical |
| 29 | NO | F | 31 | RT+LE | None | Dixi Medical |
| 30 | NO | M | 23 | LTLE | None | Dixi Medical |
| 31 | NO | F | 29 | LTLE | Left fronto-insular atrophy, left HS | Dixi Medical |
| 32 | NO | M | 44 | RTLE | None | Dixi Medical |
| 33 | NO | F | 27 | LTLE | Left superior temporal gyrus FCD | Dixi Medical |
| 34 | NO | F | 37 | LTLE | None | Dixi Medical |

**Table S1.** Demographic and clinical characteristics of the 34 recruited patients. Five patients (YES in the Excluded column) were excluded because of poor EEG recording quality, leaving 29 patients in the analyzed cohort. Abbreviations: M: male; F: female; R: right; L: left; B: bilateral (when used as a prefix to indicate laterality); TLE: temporal lobe epilepsy; PLE: parietal lobe epilepsy; FLE: frontal lobe epilepsy; OLE: occipital lobe epilepsy; T+LE: temporal-plus epilepsy; FCD: focal cortical dysplasia; HS: hippocampal sclerosis; PNH: periventricular nodular heterotopia. AD-Tech: 6–10 contacts of 2.4 mm length, 1.1 mm diameter and 2.6 mm inter-contact distance. Dixi Medical: 10–16 contacts of 2 mm length, 0.8 mm diameter and 1.5 mm inter-contact distance.

| Frequency band | ESI method | Estimated correlation [95% CI] |
|---|---|---|
| **broadband** | eLORETA | 0.090 [0.081, 0.100] |
| | sLORETA | 0.096 [0.087, 0.106] |
| | wMNE | 0.080 [0.072, 0.089] |
| | LCMV | 0.075 [0.067, 0.083] |
| **delta** | eLORETA | 0.117 [0.106, 0.130] |
| | sLORETA | 0.126 [0.114, 0.139] |
| | wMNE | 0.104 [0.094, 0.115] |
| | LCMV | 0.100 [0.091, 0.111] |
| **theta** | eLORETA | 0.113 [0.102, 0.124] |
| | sLORETA | 0.117 [0.106, 0.129] |
| | wMNE | 0.103 [0.094, 0.114] |
| | LCMV | 0.082 [0.074, 0.091] |
| **alpha** | eLORETA | 0.105 [0.095, 0.116] |
| | sLORETA | 0.109 [0.098, 0.120] |
| | wMNE | 0.100 [0.090, 0.111] |
| | LCMV | 0.077 [0.070, 0.086] |
| **beta** | eLORETA | 0.065 [0.059, 0.072] |
| | sLORETA | 0.068 [0.061, 0.076] |
| | wMNE | 0.059 [0.053, 0.066] |
| | LCMV | 0.063 [0.056, 0.070] |

**Table S2**. Estimated absolute iEEG-virtual iEEG correlations, with 95% confidence intervals, for each frequency band × ESI method combination, from the GLME described in Section 2.5.1.

| ESI method | Frequency band 1 | Frequency band 2 | Difference (band 1 — band 2) | *p*-value |
|---|---|---|---|---|
| **eLORETA** | broadband | delta | −0.027 | < 0.001 |
| | broadband | theta | −0.023 | < 0.001 |
| | broadband | alpha | −0.015 | < 0.001 |
| | broadband | beta | 0.025 | < 0.001 |
| | delta | theta | 0.005 | 0.589 (ns) |
| | delta | alpha | 0.013 | < 0.001 |
| | delta | beta | 0.052 | < 0.001 |
| | theta | alpha | 0.008 | 0.008 |
| | theta | beta | 0.048 | < 0.001 |
| | alpha | beta | 0.040 | < 0.001 |
| **sLORETA** | broadband | delta | −0.030 | < 0.001 |
| | broadband | theta | −0.021 | < 0.001 |
| | broadband | alpha | −0.013 | < 0.001 |
| | broadband | beta | 0.028 | < 0.001 |
| | delta | theta | 0.009 | 0.007 |
| | delta | alpha | 0.017 | < 0.001 |
| | delta | beta | 0.058 | < 0.001 |
| | theta | alpha | 0.008 | 0.010 |
| | theta | beta | 0.049 | < 0.001 |
| | alpha | beta | 0.040 | < 0.001 |
| **wMNE** | broadband | delta | −0.024 | < 0.001 |
| | broadband | theta | −0.024 | < 0.001 |
| | broadband | alpha | −0.020 | < 0.001 |
| | broadband | beta | 0.020 | < 0.001 |
| | delta | theta | 0.000 | 1.000 (ns) |
| | delta | alpha | 0.004 | 0.803 (ns) |
| | delta | beta | 0.045 | < 0.001 |
| | theta | alpha | 0.003 | 1.000 (ns) |
| | theta | beta | 0.044 | < 0.001 |
| | alpha | beta | 0.041 | < 0.001 |
| **LCMV** | broadband | delta | −0.026 | < 0.001 |
| | broadband | theta | −0.008 | < 0.001 |
| | broadband | alpha | −0.003 | 1.000 (ns) |
| | broadband | beta | 0.012 | < 0.001 |
| | delta | theta | 0.018 | < 0.001 |
| | delta | alpha | 0.023 | < 0.001 |
| | delta | beta | 0.038 | < 0.001 |
| | theta | alpha | 0.005 | 0.035 |
| | theta | beta | 0.020 | < 0.001 |
| | alpha | beta | 0.015 | < 0.001 |

**Table S3**. Pairwise comparisons of frequency bands within each ESI method for the iEEG-virtual iEEG correlation. Negative differences indicate a lower correlation for band 1. All *p*-values are Bonferroni-corrected; ns: not significant.

| Frequency band | ESI method 1 | ESI method 2 | Difference (method 1 – method 2) | *p*-value |
|---|---|---|---|---|
| **broadband** | eLORETA | sLORETA | −0.006 | 0.020 |
| | eLORETA | wMNE | 0.010 | < 0.001 |
| | eLORETA | LCMV | 0.015 | < 0.001 |
| | sLORETA | wMNE | 0.016 | < 0.001 |
| | sLORETA | LCMV | 0.021 | < 0.001 |
| | wMNE | LCMV | 0.005 | 0.012 |
| **delta** | eLORETA | sLORETA | −0.008 | 0.010 |
| | eLORETA | wMNE | 0.013 | < 0.001 |
| | eLORETA | LCMV | 0.017 | < 0.001 |
| | sLORETA | wMNE | 0.022 | < 0.001 |
| | sLORETA | LCMV | 0.025 | < 0.001 |
| | wMNE | LCMV | 0.004 | 0.690 (ns) |
| **theta** | eLORETA | sLORETA | −0.004 | 0.620 (ns) |
| | eLORETA | wMNE | 0.009 | < 0.001 |
| | eLORETA | LCMV | 0.030 | < 0.001 |
| | sLORETA | wMNE | 0.013 | < 0.001 |
| | sLORETA | LCMV | 0.034 | < 0.001 |
| | wMNE | LCMV | 0.021 | < 0.001 |
| **alpha** | eLORETA | sLORETA | −0.004 | 0.540 (ns) |
| | eLORETA | wMNE | 0.005 | 0.240 (ns) |
| | eLORETA | LCMV | 0.027 | < 0.001 |
| | sLORETA | wMNE | 0.009 | 0.001 |
| | sLORETA | LCMV | 0.031 | < 0.001 |
| | wMNE | LCMV | 0.023 | < 0.001 |
| **beta** | eLORETA | sLORETA | −0.003 | 0.196 (ns) |
| | eLORETA | wMNE | 0.006 | < 0.001 |
| | eLORETA | LCMV | 0.002 | 0.485 (ns) |
| | sLORETA | wMNE | 0.009 | < 0.001 |
| | sLORETA | LCMV | 0.006 | < 0.001 |
| | wMNE | LCMV | −0.003 | 0.082 (ns) |

**Table S4**. Pairwise comparisons of ESI methods within each frequency band for the iEEG-virtual iEEG correlation. Negative differences indicate a lower correlation for method 1. All *p*-values are Bonferroni-corrected; ns: not significant.

| Frequency band | Estimated PSF-PLE (cm) [95% CI] |
|---|---|
| broadband | 2.986 [2.919, 3.054] |
| delta | 3.464 [3.386, 3.543] |
| theta | 3.029 [2.961, 3.099] |
| alpha | 3.048 [2.980, 3.118] |
| beta | 2.922 [2.857, 2.989] |

**Table S5**. Estimated PSF-PLE per frequency band, with 95% confidence intervals, averaged across ESI methods (the frequency band × ESI method interaction was not significant).

| Frequency band 1 | Frequency band 2 | Difference (frequency 1 − frequency 2, cm) | *p*-value |
|---|---|---|---|
| broadband | delta | −0.478 | < 0.001 |
| broadband | theta | −0.043 | 1.000 (ns) |
| broadband | alpha | −0.062 | 0.309 (ns) |
| broadband | beta | 0.064 | 0.235 (ns) |
| delta | theta | 0.435 | < 0.001 |
| delta | alpha | 0.416 | < 0.001 |
| delta | beta | 0.541 | < 0.001 |
| theta | alpha | −0.019 | 1.000 (ns) |
| theta | beta | 0.107 | 0.002 |
| alpha | beta | 0.126 | < 0.001 |

**Table S6.** Pairwise frequency band comparisons for PSF-PLE (averaged across ESI methods).

| Metric | ESI method | Frequency band 1 | Frequency band 2 | Difference (frequency 1 − frequency 2, cm) | *p*-value |
|---|---|---|---|---|---|
| **PSF-SD** | **eLORETA** | broadband | delta | −0.145 | < 0.001 |
| | | broadband | theta | −0.153 | < 0.001 |
| | | broadband | alpha | −0.155 | < 0.001 |
| | | broadband | beta | 0.068 | 0.049 |
| | | delta | theta | −0.009 | 1.000 (ns) |
| | | delta | alpha | −0.011 | 1.000 (ns) |
| | | delta | beta | 0.213 | < 0.001 |
| | | theta | alpha | −0.002 | 1.000 (ns) |
| | | theta | beta | 0.221 | < 0.001 |
| | | alpha | beta | 0.223 | < 0.001 |
| | **sLORETA** | broadband | delta | −0.155 | < 0.001 |
| | | broadband | theta | −0.136 | < 0.001 |
| | | broadband | alpha | −0.146 | < 0.001 |
| | | broadband | beta | 0.043 | 0.850 (ns) |
| | | delta | theta | 0.018 | 1.000 (ns) |
| | | delta | alpha | 0.008 | 1.000 (ns) |
| | | delta | beta | 0.197 | < 0.001 |
| | | theta | alpha | −0.010 | 1.000 (ns) |
| | | theta | beta | 0.179 | < 0.001 |
| | | alpha | beta | 0.189 | < 0.001 |
| | **wMNE** | broadband | delta | −0.152 | < 0.001 |
| | | broadband | theta | −0.070 | 0.083 (ns) |
| | | broadband | alpha | −0.040 | 1.000 (ns) |
| | | broadband | beta | 0.176 | < 0.001 |
| | | delta | theta | 0.083 | 0.021 |
| | | delta | alpha | 0.112 | < 0.001 |
| | | delta | beta | 0.328 | < 0.001 |
| | | theta | alpha | 0.029 | 1.000 (ns) |
| | | theta | beta | 0.245 | < 0.001 |
| | | alpha | beta | 0.216 | < 0.001 |
| | **LCMV** | broadband | delta | −0.091 | 0.004 |
| | | broadband | theta | −0.078 | 0.023 |
| | | broadband | alpha | −0.133 | < 0.001 |
| | | broadband | beta | −0.125 | < 0.001 |
| | | delta | theta | 0.013 | 1.000 (ns) |
| | | delta | alpha | −0.042 | 1.000 (ns) |
| | | delta | beta | −0.033 | 1.000 (ns) |
| | | theta | alpha | −0.055 | 0.357 (ns) |
| | | theta | beta | −0.046 | 0.740 (ns) |
| | | alpha | beta | 0.008 | 1.000 (ns) |
| **CTF-PLE** | **eLORETA** | broadband | delta | −0.421 | < 0.001 |
| | | broadband | theta | −0.159 | 0.009 |
| | | broadband | alpha | −0.249 | < 0.001 |
| | | broadband | beta | −0.096 | 0.425 (ns) |
| | | delta | theta | 0.262 | < 0.001 |
| | | delta | alpha | 0.172 | 0.012 |
| | | delta | beta | 0.324 | < 0.001 |
| | | theta | alpha | −0.090 | 0.740 (ns) |
| | | theta | beta | 0.063 | 1.000 (ns) |
| | | alpha | beta | 0.153 | 0.022 |
| | **sLORETA** | broadband | delta | −0.504 | < 0.001 |
| | | broadband | theta | −0.198 | < 0.001 |
| | | broadband | alpha | −0.279 | < 0.001 |
| | | broadband | beta | −0.124 | 0.112 (ns) |
| | | delta | theta | 0.306 | < 0.001 |
| | | delta | alpha | 0.225 | < 0.001 |
| | | delta | beta | 0.380 | < 0.001 |
| | | theta | alpha | −0.081 | 1.000 (ns) |
| | | theta | beta | 0.074 | 1.000 (ns) |
| | | alpha | beta | 0.155 | 0.026 |
| | **wMNE** | broadband | delta | −0.418 | < 0.001 |

| Metric | ESI method | Frequency band 1 | Frequency band 2 | Difference (frequency 1 − frequency 2, cm) | *p*-value |
|---|---|---|---|---|---|
| | | broadband | theta | −0.233 | < 0.001 |
| | | broadband | alpha | −0.358 | < 0.001 |
| | | broadband | beta | −0.044 | 1.000 (ns) |
| | | delta | theta | 0.185 | 0.008 |
| | | delta | alpha | 0.060 | 1.000 (ns) |
| | | delta | beta | 0.374 | < 0.001 |
| | | theta | alpha | −0.125 | 0.216 (ns) |
| | | theta | beta | 0.188 | 0.002 |
| | | alpha | beta | 0.314 | < 0.001 |
| | **LCMV** | broadband | delta | −0.271 | < 0.001 |
| | | broadband | theta | −0.073 | 1.000 (ns) |
| | | broadband | alpha | −0.152 | 0.018 |
| | | broadband | beta | −0.244 | < 0.001 |
| | | delta | theta | 0.198 | < 0.001 |
| | | delta | alpha | 0.119 | 0.212 (ns) |
| | | delta | beta | 0.027 | 1.000 (ns) |
| | | theta | alpha | −0.079 | 1.000 (ns) |
| | | theta | beta | −0.171 | 0.007 |
| | | alpha | beta | −0.092 | 0.738 (ns) |
| **CTF-SD** | **eLORETA** | broadband | delta | −0.250 | < 0.001 |
| | | broadband | theta | −0.050 | 0.297 (ns) |
| | | broadband | alpha | −0.110 | < 0.001 |
| | | broadband | beta | −0.206 | < 0.001 |
| | | delta | theta | 0.200 | < 0.001 |
| | | delta | alpha | 0.141 | < 0.001 |
| | | delta | beta | 0.044 | 0.698 (ns) |
| | | theta | alpha | −0.059 | 0.114 (ns) |
| | | theta | beta | −0.156 | < 0.001 |
| | | alpha | beta | −0.097 | < 0.001 |
| | **sLORETA** | broadband | delta | −0.251 | < 0.001 |
| | | broadband | theta | −0.040 | 0.863 (ns) |
| | | broadband | alpha | −0.116 | < 0.001 |
| | | broadband | beta | −0.279 | < 0.001 |
| | | delta | theta | 0.211 | < 0.001 |
| | | delta | alpha | 0.135 | < 0.001 |
| | | delta | beta | −0.028 | 1.000 (ns) |
| | | theta | alpha | −0.076 | 0.012 |
| | | theta | beta | −0.239 | < 0.001 |
| | | alpha | beta | −0.163 | < 0.001 |
| | **wMNE** | broadband | delta | −0.221 | < 0.001 |
| | | broadband | theta | −0.061 | 0.099 (ns) |
| | | broadband | alpha | −0.155 | < 0.001 |
| | | broadband | beta | −0.211 | < 0.001 |
| | | delta | theta | 0.161 | < 0.001 |
| | | delta | alpha | 0.066 | 0.065 (ns) |
| | | delta | beta | 0.011 | 1.000 (ns) |
| | | theta | alpha | −0.094 | < 0.001 |
| | | theta | beta | −0.150 | < 0.001 |
| | | alpha | beta | −0.056 | 0.220 (ns) |
| | **LCMV** | broadband | delta | −0.160 | < 0.001 |
| | | broadband | theta | −0.015 | 1.000 (ns) |
| | | broadband | alpha | −0.094 | < 0.001 |
| | | broadband | beta | −0.402 | < 0.001 |
| | | delta | theta | 0.146 | < 0.001 |
| | | delta | alpha | 0.066 | 0.060 (ns) |
| | | delta | beta | −0.241 | < 0.001 |
| | | theta | alpha | −0.079 | 0.008 |
| | | theta | beta | −0.387 | < 0.001 |
| | | alpha | beta | −0.308 | < 0.001 |

**Table S7**. Pairwise comparisons of frequency bands within each ESI method for the reconstruction metrics.

| Metric | Frequency band | ESI method 1 | ESI method 2 | Difference (method 1 − method 2, cm) | *p*-value |
|---|---|---|---|---|---|
| **PSF-SD** | **broadband** | eLORETA | sLORETA | −0.070 | 0.026 |
| | | eLORETA | wMNE | −0.281 | < 0.001 |
| | | eLORETA | LCMV | −0.158 | < 0.001 |
| | | sLORETA | wMNE | −0.210 | < 0.001 |
| | | sLORETA | LCMV | −0.088 | 0.003 |
| | | wMNE | LCMV | 0.123 | < 0.001 |
| | **Delta** | eLORETA | sLORETA | −0.080 | 0.010 |
| | | eLORETA | wMNE | −0.288 | < 0.001 |
| | | eLORETA | LCMV | −0.105 | < 0.001 |
| | | sLORETA | wMNE | −0.208 | < 0.001 |
| | | sLORETA | LCMV | −0.024 | 1.000 (ns) |
| | | wMNE | LCMV | 0.184 | < 0.001 |
| | **Theta** | eLORETA | sLORETA | −0.053 | 0.218 (ns) |
| | | eLORETA | wMNE | −0.197 | < 0.001 |
| | | eLORETA | LCMV | −0.083 | 0.007 |
| | | sLORETA | wMNE | −0.144 | < 0.001 |
| | | sLORETA | LCMV | −0.029 | 1.000 (ns) |
| | | wMNE | LCMV | 0.114 | < 0.001 |
| | **Alpha** | eLORETA | sLORETA | −0.061 | 0.100 (ns) |
| | | eLORETA | wMNE | −0.166 | < 0.001 |
| | | eLORETA | LCMV | −0.135 | < 0.001 |
| | | sLORETA | wMNE | −0.104 | < 0.001 |
| | | sLORETA | LCMV | −0.074 | 0.026 |
| | | wMNE | LCMV | 0.030 | 1.000 (ns) |
| | **Beta** | eLORETA | sLORETA | −0.096 | < 0.001 |
| | | eLORETA | wMNE | −0.173 | < 0.001 |
| | | eLORETA | LCMV | −0.351 | < 0.001 |
| | | sLORETA | wMNE | −0.077 | 0.011 |
| | | sLORETA | LCMV | −0.255 | < 0.001 |
| | | wMNE | LCMV | −0.177 | < 0.001 |
| **CTF-PLE** | **Broadband** | eLORETA | sLORETA | −0.061 | 1.000 (ns) |
| | | eLORETA | wMNE | −0.113 | 0.108 (ns) |
| | | eLORETA | LCMV | −0.050 | 1.000 (ns) |
| | | sLORETA | wMNE | −0.051 | 1.000 (ns) |
| | | sLORETA | LCMV | 0.012 | 1.000 (ns) |
| | | wMNE | LCMV | 0.063 | 1.000 (ns) |
| | **Delta** | eLORETA | sLORETA | −0.145 | 0.058 (ns) |
| | | eLORETA | wMNE | −0.110 | 0.290 (ns) |
| | | eLORETA | LCMV | 0.100 | 0.381 (ns) |
| | | sLORETA | wMNE | 0.035 | 1.000 (ns) |
| | | sLORETA | LCMV | 0.245 | < 0.001 |
| | | wMNE | LCMV | 0.210 | < 0.001 |
| | **Theta** | eLORETA | sLORETA | −0.101 | 0.280 (ns) |
| | | eLORETA | wMNE | −0.187 | 0.002 |
| | | eLORETA | LCMV | 0.036 | 1.000 (ns) |
| | | sLORETA | wMNE | −0.086 | 0.605 (ns) |
| | | sLORETA | LCMV | 0.137 | 0.039 |
| | | wMNE | LCMV | 0.223 | < 0.001 |
| | **Alpha** | eLORETA | sLORETA | −0.092 | 0.477 (ns) |
| | | eLORETA | wMNE | −0.222 | < 0.001 |
| | | eLORETA | LCMV | 0.047 | 1.000 (ns) |
| | | sLORETA | wMNE | −0.130 | 0.102 (ns) |
| | | sLORETA | LCMV | 0.139 | 0.045 |
| | | wMNE | LCMV | 0.268 | < 0.001 |
| | **Beta** | eLORETA | sLORETA | −0.089 | 0.421 (ns) |
| | | eLORETA | wMNE | −0.061 | 1.000 (ns) |
| | | eLORETA | LCMV | −0.198 | < 0.001 |
| | | sLORETA | wMNE | 0.028 | 1.000 (ns) |
| | | sLORETA | LCMV | −0.109 | 0.203 (ns) |
| | | wMNE | LCMV | −0.137 | 0.043 |

| Metric | Frequency band | ESI method 1 | ESI method 2 | Difference (method 1 − method 2, cm) | *p*-value |
|---|---|---|---|---|---|
| **CTF-SD** | **Broadband** | eLORETA | sLORETA | −0.010 | 1.000 (ns) |
| | | eLORETA | wMNE | −0.064 | 0.033 |
| | | eLORETA | LCMV | −0.075 | 0.007 |
| | | sLORETA | wMNE | −0.054 | 0.115 (ns) |
| | | sLORETA | LCMV | −0.066 | 0.029 |
| | | wMNE | LCMV | −0.011 | 1.000 (ns) |
| | **Delta** | eLORETA | sLORETA | −0.010 | 1.000 (ns) |
| | | eLORETA | wMNE | −0.035 | 0.900 (ns) |
| | | eLORETA | LCMV | 0.015 | 1.000 (ns) |
| | | sLORETA | wMNE | −0.025 | 1.000 (ns) |
| | | sLORETA | LCMV | 0.025 | 1.000 (ns) |
| | | wMNE | LCMV | 0.050 | 0.250 (ns) |
| | **Theta** | eLORETA | sLORETA | 0.001 | 1.000 (ns) |
| | | eLORETA | wMNE | −0.075 | 0.009 |
| | | eLORETA | LCMV | −0.040 | 0.529 (ns) |
| | | sLORETA | wMNE | −0.075 | 0.008 |
| | | sLORETA | LCMV | −0.041 | 0.499 (ns) |
| | | wMNE | LCMV | 0.035 | 0.845 (ns) |
| | **Alpha** | eLORETA | sLORETA | −0.016 | 1.000 (ns) |
| | | eLORETA | wMNE | −0.110 | < 0.001 |
| | | eLORETA | LCMV | −0.060 | 0.071 (ns) |
| | | sLORETA | wMNE | −0.094 | < 0.001 |
| | | sLORETA | LCMV | −0.044 | 0.390 (ns) |
| | | wMNE | LCMV | 0.050 | 0.235 (ns) |
| | **Beta** | eLORETA | sLORETA | −0.083 | 0.004 |
| | | eLORETA | wMNE | −0.069 | 0.028 |
| | | eLORETA | LCMV | −0.271 | < 0.001 |
| | | sLORETA | wMNE | 0.014 | 1.000 (ns) |
| | | sLORETA | LCMV | −0.188 | < 0.001 |
| | | wMNE | LCMV | −0.202 | < 0.001 |

**Table S8**. Pairwise comparisons of ESI methods within each frequency band for the reconstruction metrics.

| Frequency band | ESI method | Estimated PSF-SD (cm) [95% CI] | Estimated CTF-PLE (cm) [95% CI] | Estimated CTF-SD (cm) [95% CI] |
|---|---|---|---|---|
| **broadband** | eLORETA | 3.867 [3.633, 4.116] | 2.386 [2.298, 2.476] | 4.140 [3.874, 4.425] |
| | sLORETA | 3.937 [3.699, 4.191] | 2.447 [2.357, 2.540] | 4.150 [3.883, 4.436] |
| | wMNE | 4.148 [3.897, 4.415] | 2.498 [2.406, 2.593] | 4.205 [3.934, 4.494] |
| | LCMV | 4.025 [3.781, 4.284] | 2.435 [2.346, 2.528] | 4.216 [3.944, 4.506] |
| **delta** | eLORETA | 4.011 [3.769, 4.270] | 2.806 [2.703, 2.913] | 4.391 [4.108, 4.693] |
| | sLORETA | 4.092 [3.844, 4.355] | 2.951 [2.843, 3.064] | 4.401 [4.117, 4.704] |
| | wMNE | 4.300 [4.040, 4.577] | 2.916 [2.809, 3.027] | 4.426 [4.141, 4.730] |
| | LCMV | 4.116 [3.867, 4.381] | 2.707 [2.607, 2.810] | 4.376 [4.094, 4.677] |
| **theta** | eLORETA | 4.020 [3.777, 4.279] | 2.545 [2.451, 2.641] | 4.191 [3.921, 4.479] |
| | sLORETA | 4.074 [3.827, 4.336] | 2.645 [2.548, 2.746] | 4.190 [3.920, 4.478] |
| | wMNE | 4.217 [3.962, 4.489] | 2.731 [2.631, 2.835] | 4.265 [3.991, 4.559] |
| | LCMV | 4.103 [3.855, 4.367] | 2.509 [2.417, 2.604] | 4.231 [3.958, 4.522] |
| **alpha** | eLORETA | 4.022 [3.779, 4.281] | 2.635 [2.538, 2.735] | 4.250 [3.976, 4.542] |
| | sLORETA | 4.083 [3.836, 4.346] | 2.726 [2.626, 2.830] | 4.266 [3.991, 4.559] |
| | wMNE | 4.188 [3.934, 4.458] | 2.856 [2.751, 2.965] | 4.360 [4.079, 4.659] |
| | LCMV | 4.158 [3.906, 4.425] | 2.588 [2.493, 2.686] | 4.310 [4.032, 4.606] |
| **beta** | eLORETA | 3.799 [3.569, 4.043] | 2.482 [2.391, 2.576] | 4.347 [4.067, 4.646] |
| | sLORETA | 3.895 [3.659, 4.145] | 2.571 [2.477, 2.669] | 4.429 [4.144, 4.734] |
| | wMNE | 3.972 [3.732, 4.228] | 2.543 [2.449, 2.639] | 4.415 [4.131, 4.719] |
| | LCMV | 4.149 [3.898, 4.417] | 2.680 [2.581, 2.782] | 4.618 [4.320, 4.935] |

**Table S9**. Estimated reconstruction metrics for frequency band × ESI method combinations with 95% confidence intervals.

| Frequency band | 1st | 2nd | 3rd | 4th |
|---|---|---|---|---|
| ***PSF-metrics*** | | | | |
| broadband | eLORETA[a)] | sLORETA[a)] | wMNE[b)] | LCMV[b)] |
| delta | eLORETA | sLORETA | LCMV | wMNE |
| theta | eLORETA | sLORETA | LCMV | wMNE |
| alpha | sLORETA | eLORETA | wMNE | LCMV |
| beta | eLORETA | sLORETA | wMNE | LCMV |
| ***CTF-metrics*** | | | | |
| broadband | sLORETA | eLORETA | wMNE | LCMV |
| delta | sLORETA | eLORETA | LCMV | wMNE |
| theta | eLORETA | sLORETA | LCMV | wMNE |
| alpha | eLORETA | sLORETA | LCMV | wMNE |
| beta | eLORETA | sLORETA | wMNE | LCMV |

**Table S10**. Ranking of the four ESI methods (eLORETA, sLORETA, wMNE, LCMV), combining iEEG-virtual iEEG correlation, PLE and SD, from both PSFs and CTFs. [a)] Tied for first place; [b)] tied for third place.

| ESI method | Metric | Frequency band | Fixed effects [95% CI] | *p*-value | R² |
|---|---|---|---|---|---|
| **eLORETA** | **PSF-PLE** | broadband | −0.228 [−0.253, −0.202] | < 0.001 | 0.415 |
| | | delta | −0.200 [−0.219, −0.180] | < 0.001 | 0.438 |
| | | theta | −0.234 [−0.265, −0.203] | < 0.001 | 0.452 |
| | | alpha | −0.227 [−0.258, −0.196] | < 0.001 | 0.344 |
| | | beta | −0.258 [−0.291, −0.224] | < 0.001 | 0.414 |
| | **PSF-SD** | broadband | −0.259 [−0.321, −0.196] | < 0.001 | 0.217 |
| | | delta | −0.249 [−0.297, −0.201] | < 0.001 | 0.386 |
| | | theta | −0.267 [−0.323, −0.211] | < 0.001 | 0.281 |
| | | alpha | −0.205 [−0.278, −0.131] | < 0.001 | 0.237 |
| | | beta | −0.274 [−0.359, −0.190] | < 0.001 | 0.283 |
| **sLORETA** | **PSF-PLE** | broadband | −0.236 [−0.265, −0.207] | < 0.001 | 0.393 |
| | | delta | −0.197 [−0.221, −0.173] | < 0.001 | 0.464 |
| | | theta | −0.235 [−0.264, −0.207] | < 0.001 | 0.427 |
| | | alpha | −0.237 [−0.269, −0.205] | < 0.001 | 0.379 |
| | | beta | −0.252 [−0.283, −0.222] | < 0.001 | 0.413 |
| | **PSF-SD** | broadband | −0.248 [−0.308, −0.187] | < 0.001 | 0.226 |
| | | delta | −0.201 [−0.263, −0.139] | < 0.001 | 0.395 |
| | | theta | −0.253 [−0.312, −0.195] | < 0.001 | 0.263 |
| | | alpha | −0.176 [−0.245, −0.107] | < 0.001 | 0.219 |
| | | beta | −0.215 [−0.283, −0.147] | < 0.001 | 0.259 |
| **wMNE** | **PSF-PLE** | broadband | −0.203 [−0.233, −0.173] | < 0.001 | 0.354 |
| | | delta | −0.168 [−0.188, −0.147] | < 0.001 | 0.432 |
| | | theta | −0.215 [−0.250, −0.180] | < 0.001 | 0.397 |
| | | alpha | −0.224 [−0.246, −0.202] | < 0.001 | 0.354 |
| | | beta | −0.249 [−0.280, −0.218] | < 0.001 | 0.395 |
| | **PSF-SD** | broadband | −0.237 [−0.310, −0.164] | < 0.001 | 0.214 |
| | | delta | −0.233 [−0.329, −0.138] | < 0.001 | 0.329 |
| | | theta | −0.248 [−0.306, −0.190] | < 0.001 | 0.235 |
| | | alpha | −0.170 [−0.255, −0.085] | < 0.001 | 0.211 |
| | | beta | −0.260 [−0.333, −0.187] | < 0.001 | 0.26 |
| **LCMV** | **PSF-PLE** | broadband | −0.261 [−0.295, −0.226] | < 0.001 | 0.421 |
| | | delta | −0.221 [−0.249, −0.193] | < 0.001 | 0.38 |
| | | theta | −0.244 [−0.270, −0.219] | < 0.001 | 0.425 |
| | | alpha | −0.255 [−0.287, −0.222] | < 0.001 | 0.387 |
| | | beta | −0.222 [−0.256, −0.189] | < 0.001 | 0.42 |
| | **PSF-SD** | broadband | −0.217 [−0.289, −0.145] | < 0.001 | 0.197 |
| | | delta | −0.193 [−0.243, −0.143] | < 0.001 | 0.156 |
| | | theta | −0.225 [−0.309, −0.141] | < 0.001 | 0.265 |
| | | alpha | −0.220 [−0.293, −0.147] | < 0.001 | 0.177 |
| | | beta | −0.097 [−0.163, −0.031] | 0.039 | 0.262 |
| **eLORETA** | **CTF-PLE** | broadband | −0.185 [−0.217, −0.154] | < 0.001 | 0.297 |
| | | delta | −0.165 [−0.195, −0.134] | < 0.001 | 0.366 |
| | | theta | −0.216 [−0.253, −0.179] | < 0.001 | 0.364 |
| | | alpha | −0.202 [−0.234, −0.171] | < 0.001 | 0.28 |
| | | beta | −0.267 [−0.312, −0.221] | < 0.001 | 0.396 |
| | **CTF-SD** | broadband | −0.166 [−0.252, −0.080] | 0.002 | 0.228 |
| | | delta | 0.025 [−0.091, 0.142] | 1.000 | 0.32 |
| | | theta | −0.134 [−0.196, −0.073] | < 0.001 | 0.206 |
| | | alpha | −0.225 [−0.317, −0.134] | < 0.001 | 0.215 |
| | | beta | −0.299 [−0.380, −0.217] | < 0.001 | 0.277 |
| **sLORETA** | **CTF-PLE** | broadband | −0.182 [−0.211, −0.154] | < 0.001 | 0.322 |
| | | delta | −0.150 [−0.176, −0.124] | < 0.001 | 0.398 |
| | | theta | −0.210 [−0.244, −0.177] | < 0.001 | 0.356 |
| | | alpha | −0.189 [−0.221, −0.156] | < 0.001 | 0.282 |
| | | beta | −0.252 [−0.295, −0.209] | < 0.001 | 0.405 |
| | **CTF-SD** | broadband | −0.144 [−0.230, −0.058] | 0.011 | 0.214 |
| | | delta | 0.047 [−0.065, 0.160] | 1.000 | 0.347 |
| | | theta | −0.130 [−0.180, −0.081] | < 0.001 | 0.209 |
| | | alpha | −0.204 [−0.304, −0.104] | < 0.001 | 0.223 |
| | | beta | −0.260 [−0.346, −0.174] | < 0.001 | 0.255 |
| **wMNE** | **CTF-PLE** | broadband | −0.201 [−0.235, −0.167] | < 0.001 | 0.334 |
| | | delta | −0.156 [−0.194, −0.118] | < 0.001 | 0.367 |

| ESI method | Metric | Frequency band | Fixed effects [95% CI] | *p*-value | $R^2$ |
|---|---|---|---|---|---|
| | | theta | −0.200 [−0.235, −0.165] | < 0.001 | 0.362 |
| | | alpha | −0.186 [−0.217, −0.154] | < 0.001 | 0.299 |
| | | beta | −0.248 [−0.285, −0.211] | < 0.001 | 0.356 |
| | **CTF-SD** | broadband | −0.172 [−0.245, −0.098] | < 0.001 | 0.181 |
| | | delta | −0.045 [−0.151, 0.061] | 1.000 | 0.299 |
| | | theta | −0.160 [−0.216, −0.104] | < 0.001 | 0.189 |
| | | alpha | −0.268 [−0.377, −0.160] | < 0.001 | 0.259 |
| | | beta | −0.294 [−0.368, −0.220] | < 0.001 | 0.261 |
| **LCMV** | **CTF-PLE** | broadband | −0.189 [−0.220, −0.157] | < 0.001 | 0.283 |
| | | delta | −0.163 [−0.191, −0.136] | < 0.001 | 0.33 |
| | | theta | −0.196 [−0.231, −0.162] | < 0.001 | 0.354 |
| | | alpha | −0.215 [−0.248, −0.182] | < 0.001 | 0.356 |
| | | beta | −0.243 [−0.283, −0.203] | < 0.001 | 0.424 |
| | **CTF-SD** | broadband | −0.270 [−0.339, −0.201] | < 0.001 | 0.179 |
| | | delta | −0.146 [−0.209, −0.082] | < 0.001 | 0.124 |
| | | theta | −0.194 [−0.259, −0.129] | < 0.001 | 0.226 |
| | | alpha | −0.248 [−0.302, −0.193] | < 0.001 | 0.175 |
| | | beta | −0.207 [−0.284, −0.131] | < 0.001 | 0.297 |

**Table S11**. Fixed-effect slopes (β) from GLME models assessing the relationship between PSF and CTF reconstruction metrics and iEEG-virtual iEEG correlation, across ESI methods and frequency bands.

| ESI method | Metric | Frequency band | Fixed effects [95% CI] | *p*-value | R² |
|---|---|---|---|---|---|
| **eLORETA** | Electrode depth | broadband | −0.175 [−0.245, −0.106] | <0.001 | 0.247 |
| | Electrode depth | delta | −0.102 [−0.165, −0.040] | 0.014 | 0.347 |
| | Electrode depth | theta | −0.182 [−0.253, −0.112] | <0.001 | 0.266 |
| | Electrode depth | alpha | −0.299 [−0.364, −0.235] | <0.001 | 0.247 |
| | Electrode depth | beta | −0.377 [−0.444, −0.309] | <0.001 | 0.323 |
| | Spike rate | broadband | 0.034 [0.023, 0.046] | <0.001 | 0.19 |
| | Spike rate | delta | 0.046 [0.033, 0.058] | <0.001 | 0.34 |
| | Spike rate | theta | 0.026 [0.012, 0.040] | 0.004 | 0.237 |
| | Spike rate | alpha | 0.007 [−0.007, 0.022] | 1.000 | 0.149 |
| | Spike rate | beta | 0.009 [−0.011, 0.029] | 1.000 | 0.225 |
| **sLORETA** | Electrode depth | broadband | −0.191 [−0.263, −0.120] | <0.001 | 0.272 |
| | Electrode depth | delta | −0.125 [−0.191, −0.060] | 0.002 | 0.37 |
| | Electrode depth | theta | −0.190 [−0.261, −0.120] | <0.001 | 0.276 |
| | Electrode depth | alpha | −0.310 [−0.374, −0.246] | <0.001 | 0.256 |
| | Electrode depth | beta | −0.393 [−0.462, −0.325] | <0.001 | 0.342 |
| | Spike rate | broadband | 0.033 [0.021, 0.046] | <0.001 | 0.2 |
| | Spike rate | delta | 0.042 [0.029, 0.055] | <0.001 | 0.356 |
| | Spike rate | theta | 0.025 [0.012, 0.038] | 0.002 | 0.245 |
| | Spike rate | alpha | 0.005 [−0.009, 0.019] | 1.000 | 0.147 |
| | Spike rate | beta | 0.006 [−0.014, 0.026] | 1.000 | 0.242 |
| **wMNE** | Electrode depth | broadband | −0.250 [−0.320, −0.180] | <0.001 | 0.261 |
| | Electrode depth | delta | −0.183 [−0.254, −0.113] | <0.001 | 0.339 |
| | Electrode depth | theta | −0.229 [−0.300, −0.158] | <0.001 | 0.258 |
| | Electrode depth | alpha | −0.330 [−0.394, −0.266] | <0.001 | 0.274 |
| | Electrode depth | beta | −0.418 [−0.491, −0.345] | <0.001 | 0.354 |
| | Spike rate | broadband | 0.030 [0.017, 0.042] | <0.001 | 0.164 |
| | Spike rate | delta | 0.037 [0.025, 0.050] | <0.001 | 0.307 |
| | Spike rate | theta | 0.025 [0.012, 0.039] | 0.002 | 0.212 |
| | Spike rate | alpha | 0.003 [−0.012, 0.019] | 1.000 | 0.145 |
| | Spike rate | beta | 0.008 [−0.012, 0.027] | 1.000 | 0.215 |
| **LCMV** | Electrode depth | broadband | −0.275 [−0.335, −0.216] | <0.001 | 0.229 |
| | Electrode depth | delta | −0.235 [−0.274, −0.196] | <0.001 | 0.19 |
| | Electrode depth | theta | −0.304 [−0.355, −0.253] | <0.001 | 0.322 |
| | Electrode depth | alpha | −0.401 [−0.459, −0.344] | <0.001 | 0.323 |
| | Electrode depth | beta | −0.429 [−0.486, −0.373] | <0.001 | 0.415 |
| | Spike rate | broadband | 0.017 [0.006, 0.028] | 0.022 | 0.11 |
| | Spike rate | delta | 0.020 [0.010, 0.031] | 0.002 | 0.103 |
| | Spike rate | theta | 0.005 [−0.005, 0.015] | 1.000 | 0.196 |
| | Spike rate | alpha | −0.002 [−0.013, 0.010] | 1.000 | 0.121 |
| | Spike rate | beta | −0.007 [−0.025, 0.011] | 1.000 | 0.266 |

**Table S12**. Fixed-effect slopes on the link scale for electrode depth (in cm) and spike rate (in $10\log_{10}$(spikes/min)), with 95% confidence intervals and corrected *p*-values.

| Metric | Frequency band | $R^2$ | ROI x ESI method *p*-value |
|---|---|---|---|
| Correlation | broadband | 0.851 | < 0.001 |
| PSF-PLE | broadband | 0.66 | < 0.001 |
| PSF-SD | broadband | 0.723 | < 0.001 |
| CTF-PLE | broadband | 0.585 | < 0.001 |
| CTF-SD | broadband | 0.742 | < 0.001 |
| Correlation | delta | 0.786 | < 0.001 |
| PSF-PLE | delta | 0.667 | < 0.001 |
| PSF-SD | delta | 0.643 | < 0.001 |
| CTF-PLE | delta | 0.591 | < 0.001 |
| CTF-SD | delta | 0.745 | < 0.001 |
| Correlation | theta | 0.837 | < 0.001 |
| PSF-PLE | theta | 0.604 | < 0.001 |
| PSF-SD | theta | 0.714 | < 0.001 |
| CTF-PLE | theta | 0.593 | < 0.001 |
| CTF-SD | theta | 0.716 | < 0.001 |
| Correlation | alpha | 0.882 | < 0.001 |
| PSF-PLE | alpha | 0.664 | < 0.001 |
| PSF-SD | alpha | 0.749 | < 0.001 |
| CTF-PLE | alpha | 0.628 | < 0.001 |
| CTF-SD | alpha | 0.701 | < 0.001 |
| Correlation | beta | 0.928 | < 0.001 |
| PSF-PLE | beta | 0.758 | < 0.001 |
| PSF-SD | beta | 0.819 | < 0.001 |
| CTF-PLE | beta | 0.638 | < 0.001 |
| CTF-SD | beta | 0.802 | < 0.001 |

**Table S13**. GLME model fit (adjusted $R^2$) and ROI × method interaction, by metric and frequency band.

| Frequency band | Metric | ROI | Difference (sLORETA – eLORETA) | *p*-value |
|---|---|---|---|---|
| broadband | correlation | rostral middle frontal 3 | 0.050 | 0.002 |
| delta | correlation | rostral middle frontal 3 | 0.076 | < 0.001 |
| theta | correlation | rostral middle frontal 3 | 0.052 | 0.013 |
| alpha | CTF-PLE | pars triangularis 2 | 1.409 cm | 0.009 |
| beta | correlation | middle temporal 1 | 0.020 | 0.023 |
| beta | PSF-PLE | parahippocampal 1 | 0.995 cm | 0.011 |

**Table S14**. Pairwise comparisons between eLORETA and sLORETA within each ROI, showing only significant differences. Positive differences indicate a higher value for sLORETA. All *p*-values are FDR-corrected across ROIs.
